# Viral SSB-bound ssDNA activates the bacterial anti-phage defense system DARNA

**DOI:** 10.64898/2026.02.22.705581

**Authors:** Rugile Puteikiene, Christopher Vassallo, Arunas Silanskas, Jonas Juozapaitis, Inga Songailiene, Michael T. Laub, Giedrius Sasnauskas

## Abstract

To protect themselves against phage infection, bacteria employ diverse defense systems that are typically activated specifically upon infection. However, the mechanisms of activation and self versus non-self discrimination for most systems remain poorly understood. Here, we show that the bacterial immunity protein DARNA, once activated, cleaves a subset of host tRNAs, thereby inhibiting phage propagation. Although phages escape DARNA-mediated defense through mutations in the gene encoding single-stranded DNA-binding protein (SSB), we find that phage SSBs do not directly stimulate DARNA. Instead, DARNA is activated by single-stranded DNA presented by phage SSB, but not by the host SSB. The recognition of an endogenous nucleic acid signal promoted by a viral protein ensures that DARNA can detect and respond to a broad range of viruses while avoiding auto-immunity.

---

Bacteriophages are a major threat to bacteria. Consequently, bacteria have evolved diverse mechanisms to protect themselves against viral infections. For many anti-viral defense systems, the activity of their effector proteins can be predicted from the protein sequences, such as DNase, RNase, NADase, or membrane-disrupting domain ^1–3^. In contrast, the mechanisms underlying immune system activation upon viral infection are often difficult to infer and remain largely unknown. Importantly, the cue or trigger that activates a given defense system must be a highly specific signature of infection to prevent inappropriate activation and auto-immunity.

In some cases, defense systems are activated by individual phage structural proteins that differ from host proteins and thus provide a signature of infection. For example, Avs systems recognize the large terminase or portal protein of certain phages ^2^ while CapRel^SJ46^ recognizes the major capsid protein of some of the phages it defends against ^4,5^. In other cases, defense systems are directly triggered by phage-encoded inhibitors of other defense systems, as with CapRel^SJ46^, PrrC, and PARIS ^5–7^. There are also examples such as the Hna, AbpAB, and retron-Eco8 systems that respond to a component of the phage replication machinery, single-stranded DNA-binding protein (SSB) ^8–10^, that presumably differs enough from the host SSB that it serves as a reliable signature of infection. Recognition of infection signals, either directly leads to activation of defense systems, or it can activate second messenger pathways, in which a signal-sensing component of the defense system synthesizes small signaling molecules, thereby amplifying the signal, which is then recognized by the effector proteins. Many defense systems can, when activated, form large oligomeric assemblies, likely to enable cooperative, switch-like transitions between inactive and active states; examples include filamentous assemblies in Thoeris ^11^, CBASS ^12,13^, CalpL CRISPR-Cas auxiliary protein ^14^, as well as higher-order oligomers formed by Gabija, Sir2–HerA, and RADAR proteins ^15–18^.

In addition to protein-based triggers, some defense systems are activated by phage nucleic acids. Restriction-modification systems typically recognize and then cleave unmethylated phage DNA, while CRISPR-Cas systems recognize phage sequences matching RNA guide sequences adjacent to protospacer adjacent motifs to trigger phage DNA cleavage. The Shedu^19^ system recognizes DNA ends that arise in phage genomes but typically not in the host genome. There are hints that other systems such as Nhi ^20^ and Gabija ^21^ also recognize phage-specific nucleic acid structures, suggesting that this may be a common mechanism of defense activation.

Here, we focus on the recently discovered anti-phage defense system DARNA (ssDNA-Activated RNase), previously referred to as PD-T7-3, which functions through an abortive infection mechanism ^22^. DARNA encodes an N-terminal HEPN (Higher Eukaryotes and Prokaryotes Nucleotide-binding) RNase domain, characterized by the R_117_X_4_H_122_ catalytic motif, and a C-terminal domain of unknown function. HEPN domains, which typically dimerize to form bipartite catalytic centers at the subunit interface, are widespread in all domains of life. For example, in eukaryotes, HEPN-domain proteins play diverse roles in RNA processing and innate immunity (e.g., RNase L in mammalian cells), whereas in prokaryotes HEPN domains are found in various anti-viral systems, including ApeA, PD-T4-2, HepS CRISPR-Cas nucleases Cas13 and Csm6, as well as in TA systems such as HepTA ^2,22–28^. Overall, HEPN domain-containing proteins exhibit diverse mechanisms of regulation. They can be inhibited through protein-protein interactions (as in the PrrC tRNA-cleaving toxin) ^29^, by RNA-antitoxins (as in AbiF ^30^), or by covalent modifications (as in HepTA ^23,26^). In type III CRISPR-Cas systems, the activity of the Csm6 mRNA nuclease is activated by cyclic oligoadenylate signaling molecules produced by Cmr/Csm complexes upon phage infection ^31,32^, whereas in eukaryotes, the Las1 nuclease is regulated by its binding partner Grc3 ^33^. Upon activation, HEPN nucleases cleave a diverse set of RNA targets, including pre-rRNA (Las1 ^33^), mRNA (Csm6 ^31,32^, HepT ^26^, Cas13 ^24^), and tRNA (PrrC ^29^, HepT ^23^, Cas13 ^34^, HepS ^27^, ApeA ^28^), or deplete cyclic oligoadenylate signals ^35,36^.

We demonstrate that upon viral infection, DARNA cleaves cellular tRNAs in the anticodon loop. Although phages can escape DARNA via mutations that affect their SSB proteins, we find, surprisingly, that phage SSBs are not the key, direct trigger of DARNA. Instead, DARNA is activated by single-stranded DNA (ssDNA), with the phage SSB promoting this interaction. Cryo-EM structures reveal that ssDNA is bound simultaneously by multiple C-terminal domains present in a large, dodecameric assembly of DARNA, which represents the active form of the protein. In cells, DARNA selectively responds to ssDNA bound by phage SSB, but not by bacterial SSB, thereby ensuring that DARNA is only activated upon infection. Collectively, our results reveal a mechanism of immune activation in which a viral protein modulates the ability of an endogenous nucleic acid signal to trigger a defense system.

## Results

### Activation of DARNA is triggered by diverse phage ssDNA-binding proteins

Consistent with our prior work ^22^, we found that DARNA provided strong defense against T3, T5, and T7 phages. Cells producing DARNA from its native promoter on a low-copy plasmid showed substantial reductions in the efficiency of plating compared to an empty vector control in each case, as well as smaller plaques (Figure 1A). To understand how DARNA is activated, we screened for mutants of T3, T5, and T7 that could evade defense by DARNA. For T3 and T7, the escape mutants almost completely restored infectivity, with more modest restoration for T5 escape mutants. Whole-genome sequencing revealed that each escaper had a mutation in the gene encoding a single-stranded DNA binding protein (SSB) (Figure 1A), suggesting that SSB is necessary for activation. To test whether SSB is sufficient to trigger DARNA activation, we induced the production of T7 SSB (Gp2.5) in cells harboring DARNA and found that cells stopped growing ∼90 min post-induction (Figure 1B). In contrast, inducing T7 SSB in an empty vector control strain or inducing the escape mutant variant, SSB(S132P) in the presence of DARNA strain did not substantially affect growth (Figure 1B). Together, these results indicated that phage SSB is critical for the activation of DARNA.

**Figure 1.**
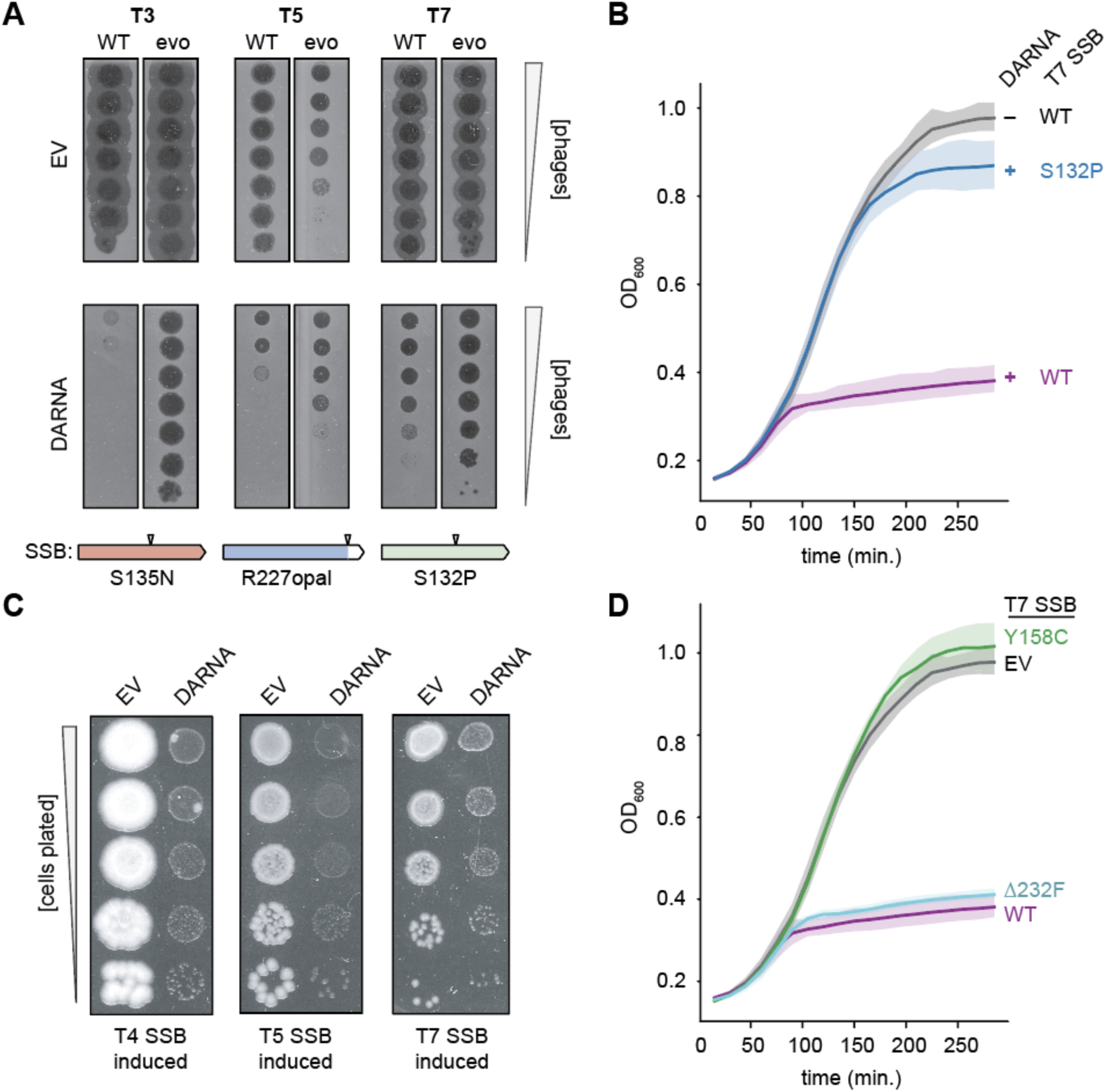
DARNA senses the activity of diverse phage single-stranded DNA binding proteins. (A) Plaque assay showing the ability of WT (ancestor) and evolved phages to form plaques on a lawn of *E. coli* with an empty vector (EV), or expressing DARNA from its native promoter. Mutations to each evolved plage’s respective SSB are shown below. (B) Measurement of bacterial growth in strains encoding DARNA (+) or EV (–) and inducing the expression of either Gp2.5 WT or the Gp2.5 S132P mutant. (C) Bacterial colony growth assay in DARNA or EV cells when inducing T7, T5 or T4 SSB proteins from pBAD. (D) Measurement of bacterial growth in strains encoding DARNA and expressing WT or mutant versions of Gp2.5.

Notably, while T3 and T7 SSBs are highly similar in structure and sequence, T5 SSB is markedly different (Figures S1A and S1B). Phage T4, which we did not generate escape mutants for, but which DARNA also defends against, encodes an SSB that is also highly divergent in sequence and structure relative to T3, T7, and T5 (Figures S1A and S1B). To test whether these disparate phage SSBs were each sufficient to activate DARNA, we induced expression of T7, T5, or T4 SSB in cells containing either DARNA or an empty vector (EV) and monitored colony formation on solid media. The induction of all three distinct SSBs impaired colony growth in cells harboring DARNA but not in cells harboring the empty vector, suggesting that all three SSBs are sufficient for activating DARNA despite their divergent sequences and structures (Figure 1C).

We next wondered which specific function of T7 SSB activates DARNA. T7 SSB protein has an N-terminal OB-fold that binds ssDNA and an acidic, unstructured C-terminal tail that promotes binding cooperativity and interaction with T7 DNA polymerase and T7 helicase/primase during replication ^37,38^. A key residue in the OB-fold is tyrosine 158 as substitutions in this residue significantly decrease ssDNA binding affinity ^39^, and deletion of the C-terminal phenylalanine, Δ232F, which prevents interaction with T7 DNA polymerase without impacting ssDNA binding ^40^. We found that co-producing SSB(Y158C) with DARNA did not suppress cell growth, whereas co-production of SSBΔ232F with DARNA produced a growth inhibition comparable to that seen when using WT SSB (Figure 1D). Together, these results suggest that DARNA detects ssDNA-binding by T7 SSB, resulting in activation of HEPN activity, host cell growth arrest, and abortive infection.

### DARNA is a promiscuous tRNA anti-codon nuclease

We next investigated the mechanism of growth arrest in cells containing active DARNA. DARNA contains a HEPN domain, often implicated in endonucleolytic cleavage of RNA. To test the hypothesis that DARNA targets RNA following phage infection, we collected cells harboring DARNA or an empty vector 12 minutes post-T7 infection, and performed RNA sequencing. We enumerated the number of read ends (normalized to the total read end count as read ends per million (REPM)) across the MG1655 genome. We then asked at which positions along the genome we obtained a high number of read ends in the DARNA containing cells compared to the control strain, with increased read ends likely arising at positions of RNA cleavage (Figure 2A). The largest signals occurred within most tRNAs (Figure 2A), with the strongest total 5′ and 3′ ends signal within *trpT*, *thrU*, *tyrT,* and *tyrV* (Figure 2A-B), but with strong signals detected in the majority of the 84 tRNAs in MG1655 (Figure 2C). For each tRNA, the 3’ end produced in cells harboring DARNA corresponded to the anti-codon loop, *e.g.*, there was a large enrichment of 3′ ends at position 34 within the anticodon loop of *trpT* (Figures 2B and 2D). For *trpT* there was a substantial drop in read coverage to the 3′ side of nucleotide 34A (Figure 2B). While the majority of tRNAs shared this pattern of increased read counts in the 3’ half of the tRNA, some showed increased read counts in the 5’ half of the tRNA or comparable read counts in each half (Figure 2C). No large differences in the number of read ends between samples was observed when we mapped reads to the T7 genome (Figure S2A). Collectively, these results indicate that DARNA, when activated by phage infection, promiscuously cleaves the anti-codon loops of most tRNAs to arrest translation and stop phage development.

**Figure 2.**
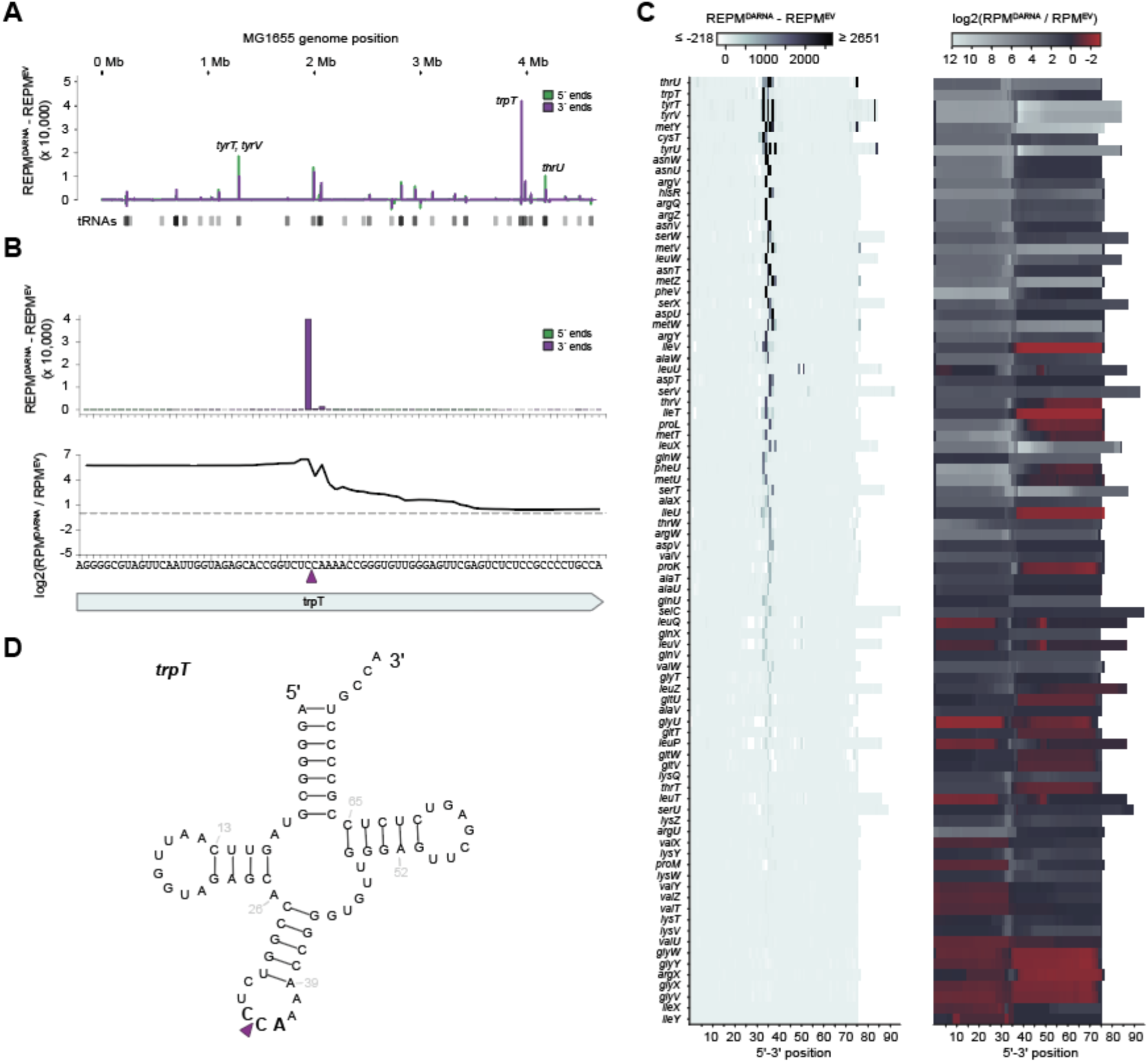
DARNA cleaves multiple tRNAs at their anticodon loops. (A) RNA-sequencing data of T7 infected empty vector cells and DARNA-encoding cells expressed as 5′ and 3′ normalized read ends (REPM) of the EV strain subtracted from the 5′ and 3′ normalized read ends of the DARNA strain at each position along the MG1655 chromosome. Positions of tRNAs are shown below and the highest scoring tRNAs are labeled (see Fig. 2C, below) (B) Top, same data as (A), but enlarged to show the *trpT* tryptophan tRNA gene. Bottom, coverage ratio at positions along the *trpT* gene. The purple arrow indicates the presumed cleavage site. (C) Left, heatmap showing the same read end metric as (A) but within 84 canonical tRNAs and scoring the sum of 5′ and 3′ ends at each position. To better show detail, the colorbar legend spans the 99.5 and 0.05 percentile of the total data. Right, coverage ratio at positions along tRNAs. (D) Sequence and schematic of *trpT* showing the site of cleavage between bases 34 and 35.

We also performed RNA sequencing on cells in which we induced the production of T7 SSB/Gp2.5 or SSB(S132P) in cells producing DARNA. Inducing WT SSB resulted in a similar cleavage profile when compared to the SSB(S132P) control, again indicating cleavage within the anti-codon loops of tRNAs (Figures S2B-S2D). These results support our conclusion that DARNA, when activated, cleaves tRNAs to arrest translation.

### tRNA cleavage is activated by ssDNA

We reconstituted tRNA cleavage in vitro using purified DARNA, phage T7 SSB/Gp2.5 and an ssDNA oligonucleotide, along with a mixture of *E. coli* tRNAs (Figure 3A). No tRNA cleavage was observed with DARNA alone or with DARNA plus T7 SSB. However, efficient tRNA cleavage was observed upon addition of ssDNA, yielding tRNA fragments approximately 30-40 nt in length. This tRNA cleavage pattern was consistent with the RNA sequencing results (Figure 2) indicating that tRNAs were predominantly cleaved within the anticodon loop. Unexpectedly, ssDNA alone stimulated tRNA cleavage by DARNA, though less efficiently compared to ssDNA together with T7 SSB. These results indicated that DARNA functions in vitro as an ssDNA-activated RNase, with SSB possibly serving to stabilize a conformation of ssDNA or to present ssDNA in a manner that promotes DARNA activation.

**Figure 3.**
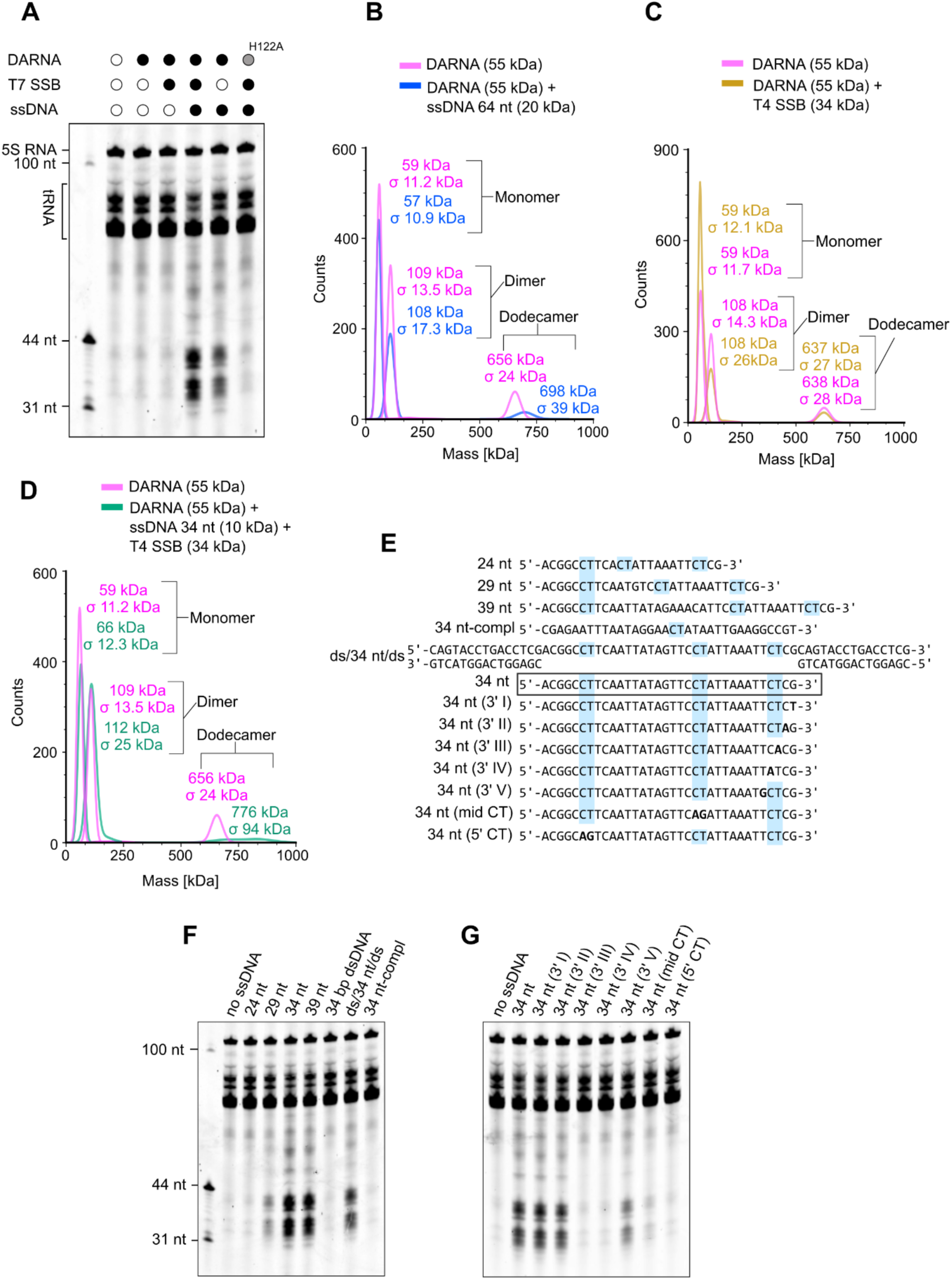
DARNA activation and interactions with ssDNA and phage SSB. (A) tRNA cleavage activity of DARNA is activated by ssDNA and phage SSB. The in vitro tRNA cleavage assays were performed using different combinations of DARNA, the DARNA catalytic center mutant H122A, the 34 nt ssDNA oligonucleotide (‘34nt’), and T7 SSB protein gp2.5. (B) DARNA interactions with ssDNA. Mass photometry particle size distributions are shown for apo DARNA (magenta) and DARNA with ssDNA (blue). DARNA concentration was 100 nM (in terms of monomer), and ssDNA concentration was 50 nM. (C) The lack of direct interactions between DARNA and bacteriophage SSB. Mass photometry particle size distributions are shown for apo DARNA (magenta) and DARNA with phage T4 SSB (yellow). DARNA and phage T4 SSB concentrations were 100 nM and 150 nM, respectively (in terms of monomer). (D) DARNA interactions with ssDNA and bacteriophage SSB. Mass photometry particle size distributions are shown for apo DARNA (magenta) and DARNA with ssDNA and phage T4 SSB (green). DARNA and phage T4 SSB concentrations were 100 nM and 150 nM, respectively (in terms of monomer), and ssDNA concentration was 50 nM. (E) Sequences of ssDNA oligonucleotides used as activators of tRNA cleavage by DARNA. The standard ‘34nt’ activator oligonucleotide is encircled. Light blue boxes highlight the ‘CT’ dinucleotide motifs. (F) tRNA cleavage assays using different single-and double-stranded DNA oligonucleotides as activators. (G) Effects of substitutions in the standard ‘34nt’ ssDNA activator DNA on tRNA cleavage. tRNA cleavage reactions in panels A, F and G contained DARNA and T7 SSB at concentrations of 0.5 μM and 5 μM, respectively (in terms of monomer), and ssDNA at 0.1 μM.

To probe the oligomeric state of DARNA, we used mass photometry, which revealed that DARNA exists in multiple states: monomeric (55 kDa); homodimeric (110 kDa), which is typical for HEPN-domain proteins; and dodecameric (660 kDa) (Figure 3B). Addition of ssDNA oligonucleotide had no effect on the molecular mass of the monomeric and dimeric forms, but increased the mass of the dodecamer, suggesting that only the higher-order oligomer interacts directly with ssDNA and thus may represent the functional form of the enzyme (Figure 3B). Addition of SSB did not alter the mass of DARNA, implying that under the experimental conditions used for mass photometry, it does not stably interact with DARNA (Figure 3C). However, addition of both SSB and ssDNA to purified DARNA protein produced particles with a broad mass distribution exceeding that of the DARNA dodecamer, indicating that the SSB protein associates with DARNA oligomers via ssDNA (Figure 3D).

To further understand how ssDNA activates DARNA, we tested a series of oligonucleotides variants (Figure 3E). The tRNA cleavage by DARNA shown in Figure 3A relied on a 34 nt ssDNA oligonucleotide ‘34nt’. DARNA was not activated by double-stranded DNA, but an ssDNA fragment flanked by duplex DNA at both ends was an efficient activator, indicating that DARNA recognition is not restricted to free ssDNA ends (Figure 3F; lanes 6, 7). Activation also required a minimum ssDNA length of approximately 30 nt (Figure 3F; lanes 2-5). Unexpectedly, the complementary strand (‘34nt-compl’) of the efficient activator ‘34nt’ failed to stimulate tRNA cleavage, suggesting that DARNA recognizes a specific sequence motif present in ‘34nt’, but absent in its complement (Figure 3F; lane 8). To test this hypothesis, we introduced single-nucleotide substitutions into the ‘34nt’ DNA, starting at the 3’ end. Mutations at the third (T) or fourth (C) position from the 3′ end, but not at the first, second or fifth positions from the 3’ end (Figures 3E and 3G), abolished activation. This finding implies that DARNA may recognize the sequence motif ‘CT’. Notably, the ‘34nt’ DNA contains three ‘CT’ motifs (two internal and the 3’-terminal subjected to mutation), whereas the non-activating ‘34nt-compl’ DNA contains only one ‘CT’ motif. Replacement of either of the two internal ‘CT’ motifs in the ‘34nt’ DNA to ‘AG’ also reduced or abolished tRNA cleavage by DARNA (Figures 3E and 3G). Taken all together, our results indicate that ssDNA containing multiple’CT’ motifs is a potent, direct activator of DARNA, with phage T7 SSB likely promoting the binding of ssDNA to DARNA to trigger its activation.

### Overall oligomeric assembly of DARNA

To uncover the structural basis of oligomeric assembly and ssDNA-mediated activation of DARNA, we determined cryo-EM structures of the protein in both apo-and nucleic acid-bound forms.

In the apo-state, the DARNA monomer forms a bent rod-shaped α-helical structure (Figures 4A and S3A). The structure consists of the N-terminal catalytic HEPN domain (aa 1-159, the closest matches revealed by DALI ^41^ search include HEPN domains of type III CRISPR-Cas Csm6 ribonucleases, e.g. PDB: 6TUG, Figure S3B). The C-terminal non-catalytic domain (aa 160-456) exhibits an overall fold reminiscent of the cohesin gatekeeper subunit Pds5, as detected by GTalign ^42^ (PDB: 5f0o, Figure S3C).

**Figure 4.**
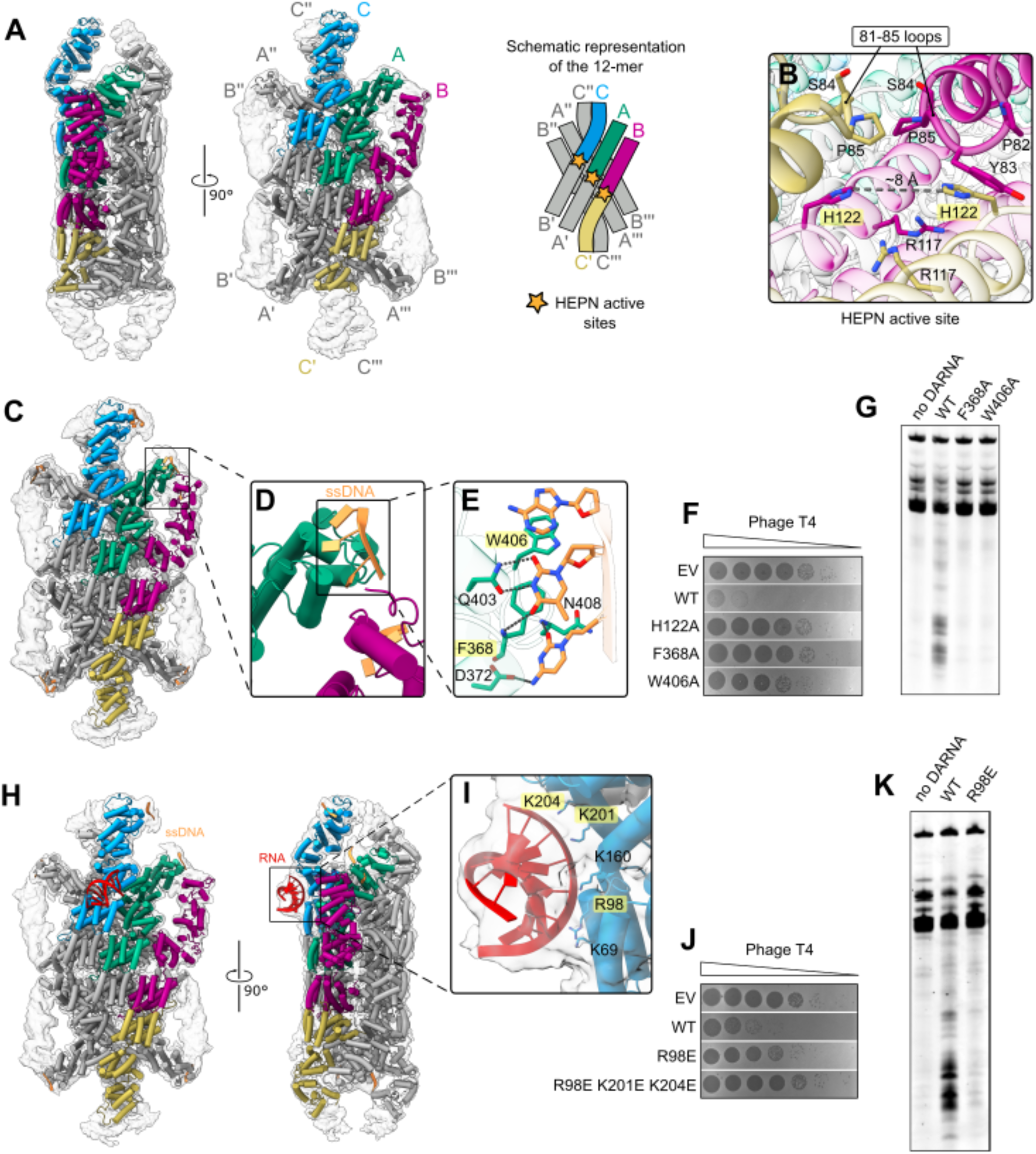
Key features of the DARNA structure. (A) Cryo-EM structure of the apo DARNA homododecamer. The unsharpened cryo-EM map and the atomic model are shown. The schematic on the right depicts the arrangement of twelve subunits—six per layer—in the DARNA complex. (B) DARNA HEPN active site located between the ‘B’ (dark purple) and ‘C’ (wheat) subunits. The 81-85 regulatory loops that occlude access to the catalytic residues H122 and R117 are shown. Substituted residues are highlighted in yellow. (C) Cryo-EM structure of DARNA bound to ssDNA. The unsharpened cryo-EM map and the atomic model are shown. A black rectangle highlights the claw-like structure formed by the C-terminal domains of subunits ‘A’ and ‘B’ that interact with ssDNA. (D) Enlarged view of the region highlighted in panel C. The black rectangle indicates the distal part of the subunit ‘A’ C-terminal domain that forms base-specific interactions with the ‘CT’ DNA sequence. (E) Detailed view of the region outlined in panel D, showing sequence-specific recognition of the ‘CT’ DNA fragment by residues of the subunit ‘A’ C-terminal domain. Substituted residues are highlighted in yellow. (F) Phage T4 plaque assay using DARNA variants with substituted residues involved in ssDNA recognition. EV - empty vector control. (G) In vitro tRNA cleavage assay using the DARNA ssDNA-recognition mutants F368A and W406A. (H) Overall structure of the DARNA homododecamer bound to ssDNA and an RNA fragment. The unsharpened cryo-EM map (transparent) and the atomic model are shown. (I) DARNA residues contacting the bound RNA in the DARNA-ssDNA-RNA complex. Substituted residues are highlighted in yellow. (J) Phage T4 plaque assay using DARNA variants carrying substitutions of the RNA-binding residues. (K) Effect of RNA binding interface substitution R98E on tRNA cleavage. tRNA cleavage reactions in panels G and K contained DARNA and T7 SSB at concentrations of 0.5 μM and 5 μM, respectively (in terms of monomer), and ssDNA at 0.1 μM.

Three elongated primary dimers, each formed through dimerization of the N-terminal HEPN domains, stack in a staggered arrangement to form a flat, diamond-shaped homohexameric structure (Figure 4A). Two such hexamers further assemble into a dodecameric, two-layered complex (Figure 4A). The overall D2 symmetry of the dodecameric assembly dictates the presence of three distinct types of the monomeric subunits, which we designate as A (central), B (lateral) and C (apical) (Figure 4A). Accordingly, two types of primary dimers are observed: A+A’ dimers at the center of the hexamer and B+C’ (or B’+C) dimers at the sides (Figure 4A).

The HEPN and C-terminal domains of the A, B and C subunits adopt nearly identical conformations (rmsd < 1 Å for the overlayed regions), differing only in the relative orientation of the two domains. The C-terminal domains of A+B and A’+B’ subunits within the same hexameric layer are positioned in close proximity, forming claw-like protrusions on the opposite sides of the hexamer. By contrast, the C-terminal domain of each C subunit is positioned against its counterpart from the opposing hexameric layer, forming claw-like protrusions at the apices of the dodecamer (Figure 4A).

DARNA assembly into the dodecameric complex is mediated by three types of protein-protein interfaces (Figure S4A):

1. HEPN domain dimerization interfaces (approx. 1000 Å^2^ for A+A’ dimers, and approx. 1200 Å^2^ for B+C’ and B’+C dimers), which are formed by the symmetrically arranged α2-α3 helices and the 121-134 loops. The surface is largely polar and is stabilized by multiple H-bonds and salt bridges.
2. Dimer-dimer interfaces (approx. 1800 Å^2^) within the hexameric layer. The mostly polar interactions are formed between the primary A-A’ and B-C’ (or B’-C) dimers. The interfaces include HEPN-HEPN contacts (e.g., helices α6-α7 of subunit A are positioned against the equivalent helices of subunit B’), HEPN-C domain contacts (e.g., helix α3 of subunit A against α10 of subunit B), and C-C domain interactions (e.g., helix α10 of subunit A against α10 of subunit B, and helix α18 contacting helices α9-α10 of subunit C).
3. Inter-hexamer interface (∼1500 Å^2^) is formed primarily by the symmetry related A-A’-C subunits from the opposite hexameric layers. The contacts include H-bonds and salt bridges that involve helices α14 and α22 of A+A’ subunits, α1, α2 and α7 of subunit C, and N-terminal regions of A+A’.

The inter-hexamer interface substitutions F6A, N33A+E347A abolished formation of dodecameric DARNA complexes (Figure S4B) and impaired DARNA activity both *in vivo* and *in vitro* (Figures S4C and S4D), indicating that dodecamer formation is required for DARNA activity.

The DARNA dodecamer contains six HEPN active sites in total (three per hexameric layer, formed between A+A’ and B+C’/B’+C dimer interfaces), all oriented toward the outer surface of the complex (Figures 4A and 4B). The catalytic histidines (H122) in each active site are separated by approximately 8 Å (Figure 4B), a distance comparable to that observed in the active conformations of Csm6 (e.g., PDB:6tug) and HepT toxin (e.g., PDB:7ae8). In DARNA, however, the catalytic centers are blocked by the loops connecting helices α4 and α5 (residues 81-85, Figures 4B and S5A). Equivalent structural elements in other HEPN nucleases are either absent (HepT), or adopt different spatial positions that do not interfere with the access to the catalytic center (Figure S4B). We therefore presume that the α4-α5 structural element may contribute to regulation of DARNA activity.

### Mechanism of ssDNA recognition

To investigate how DARNA recognizes its ssDNA activator, we prepared cryo-EM samples with the ssDNA oligonucleotide ‘34nt’, which efficiently stimulated tRNA cleavage *in vitro* (Figure 3). The cryo-EM map revealed extra density corresponding to ssDNA near the distal regions of the C-terminal domains (Figure 4C). We modeled short ssDNA fragments bound to the C-terminal domains of A, B and C subunits (Figures 4C and 4D).

The protein-DNA contacts, best resolved in the vicinity of the C-terminal domain of the A subunit (Figures 4D, 4E and 5SA), uncovered the structural basis for the observed sequence-specific nature of DARNA activation by ssDNA. The ssDNA fragment’CTA’ interacts with the protein exclusively though its bases, without direct interactions to the backbone phosphates. The cytosine forms two base-specific H-bonds with the side chain of D372 and backbone amine of N408, and a vdW contact to F368. The adjacent thymine stacks against F368, and makes three base-specific H-bonds: two with the side chain of Q403, and one with the backbone amine of F368 (Figure 4E). The third base, adenine, makes no base-specific H-bonds, but stacks against W406, and makes vdW contacts with N402 and Y405 (Figure 4E). These interactions explain the observed requirement for a ‘CT’ motif in the activating ssDNA oligonucleotides. Furthermore, the relative spatial arrangement of the ssDNA binding motifs across the adjacent C-terminal domains of the C-A-B subunits (Figure 4C) suggests that the same ssDNA fragment may simultaneously interact with multiple subunits, explaining the preference of the protein for longer ssDNA oligonucleotides containing multiple ‘CT’ sequence motifs. Substituting F368 or W406 with alanine rendered DARNA inactive both with respect to anti-phage defense activity in vivo and when reconstituted in vitro (Figures 4F and 4G).

### tRNA-bound structure

We also sought a structure of the inactive DARNA variant harboring the substitution H122A in the presence of ssDNA and an *E. coli* tRNA mix. The resulting cryo-EM map revealed density on the outer surface of the C subunit at the positively charged patch located at the N-and C-domain junction, which is consistent with a short A-form RNA fragment (Figure 4H). Although the low local resolution and small volume of this extra density precluded accurate identification and modeling of the bound RNA, it enabled us to identify the positively charged C-subunit residues that interact with the RNA backbone (Figure 4I). Substitutions at this RNA binding surface (DARNA variants R98E and R98E+K201E+K204E) rendered the protein inactive both in vivo (Figure 4J) and in vitro (Figure 4K). A similar extra density was also observed in a sample containing DARNA incubated with *E. coli* tRNA in the absence of ssDNA (Figures S5C and S5D), indicating that ssDNA activator is not required for RNA binding by the DARNA protein.

Notably, the overall DARNA protein conformation in the ssDNA-bound and ssDNA+RNA-bound complexes, including the catalytic center regions, was virtually indistinguishable from that of the apo structure, in which the catalytic centers are blocked by the 81-85 aa loop connecting helices α4 and α5. Activation of RNA cleavage likely involves a conformational change in the α4-loop-α5 region that depends on interactions with both the RNA substrate and the ssDNA activator. Consistent with this model, the active-site-occluding helices α4 and α5 also carry the RNA-binding residues K69 and R98 (Figure 4I). However, the precise molecular mechanism of DARNA activation, including the respective roles of ssDNA and RNA binding, remains to be elucidated.

### DARNA reactions in the context of bacterial SSB

For DARNA to function as a bacterial immunity system capable of self/non-self discrimination, it must not be triggered by ssDNA generated in uninfected cells, for example during DNA replication, or by bacterial SSB bound to such DNA. Consistent with this, overproducing *E. coli* SSB (EcSSB) in cells harboring DARNA did not lead to any detectable toxicity or growth defect (Figure 5A). Moreover, increasing concentrations of EcSSB inhibited rather than activated DARNA RNase activity in vitro (Figure 5B), presumably by sequestering the ssDNA activator. By contrast, phage SSB introduced into the reactions rescued tRNA cleavage in a concentration-dependent manner (Figure 5B). This finding implies that phage SSB competes with EcSSB, and that ssDNA, when bound by phage SSB rather than EcSSB, retains the ability to activate the DARNA protein. Notably, phage T4 SSB, which has higher affinity for ssDNA than T7 SSB (as determined by EMSA, Figure S6A), was a more efficient competitor to EcSSB (Figure 5B). Furthermore, T7 SSB escape variants S132P and Y158C (Figure 1A, 1D) failed to compete with EcSSB under conditions where the WT T7 protein rescued DARNA activity (Figure S6B). For the Y158C mutant this also correlated with its reduced affinity for ssDNA (Figure S6A). Collectively, our results indicate that ssDNA is the direct activator of DARNA, with this interaction promoted by phage SSB but not host SSB. Consequently, DARNA remains inactive until phage infection occurs and phage SSB in complex with ssDNA triggers activation.

**Figure 5.**
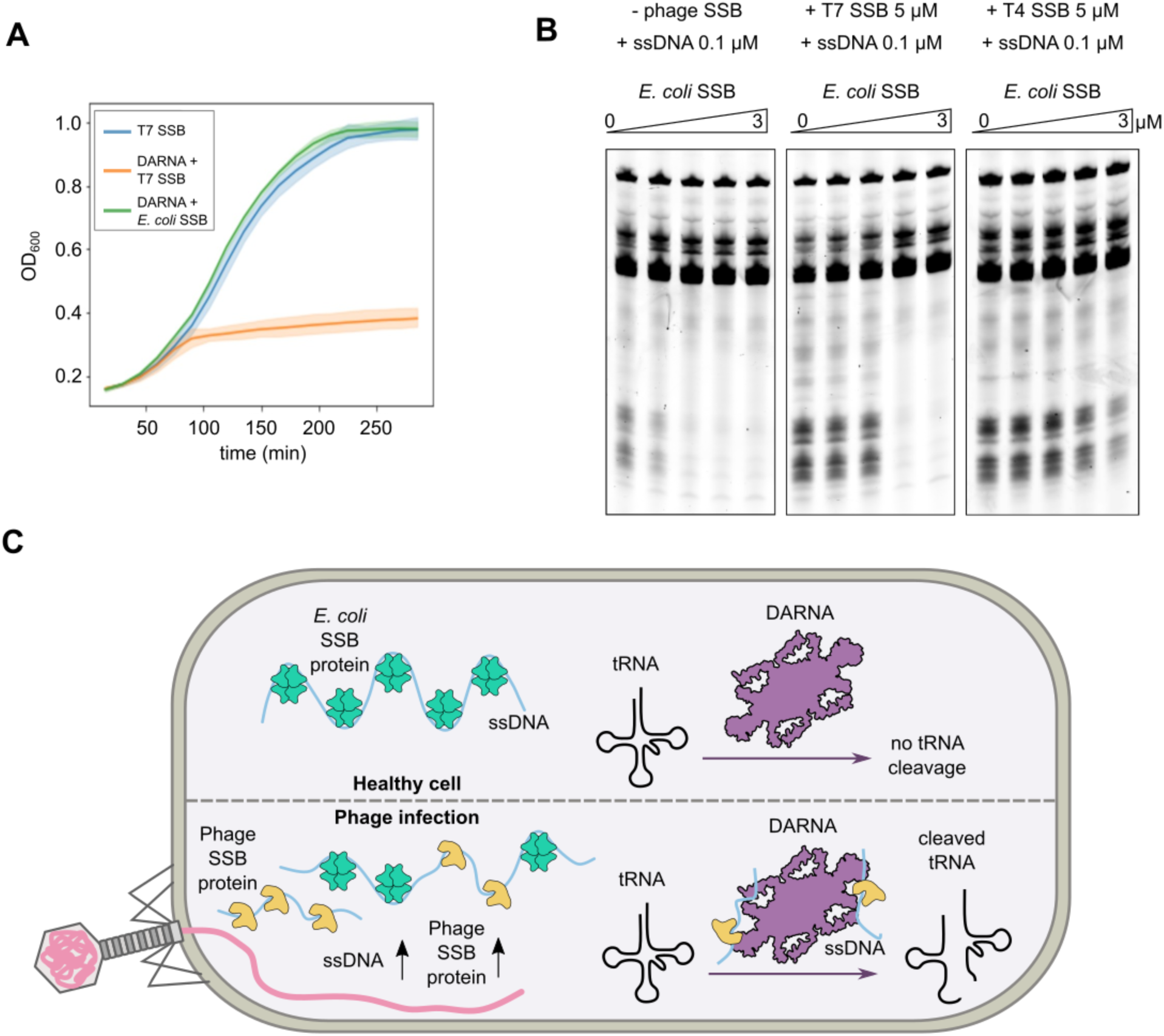
DARNA activity in the context of bacterial SSB protein. (A) Measurement of bacterial growth in strains encoding DARNA with either T7 SSB or *E. coli* SSB and strain encoding only T7 SSB (B) tRNA cleavage assays performed in the absence of phage SSB, or in the presence of phage SSB (5 µM T7 SSB or T4 SSB monomer) together with increasing concentrations of E. coli SSB (0, 0.4, 0.8, 1.6, 3.0 µM monomer). DARNA was present at a concentration of 0.5 µM (in terms of monomer) and ssDNA at 0.1 µM. (C) Proposed mechanism of DARNA action. Top: in uninfected cells, host SSB sequesters all ssDNA present in the cell, preventing activation of DARNA. Bottom: During phage infection, phage SSB competes with bacterial SSB for ssDNA fragments. ssDNA bound by phage SSB remains accessible to DARNA binding, thereby activating the tRNA cleavage activity of DARNA.

## Discussion

We characterized the structure, host tRNA cleavage activity, and activation mechanism of the previously identified anti-viral defense protein DARNA. Using cryo-EM, we found that DARNA forms a large dodecameric complex, in which the N-terminal halves of 12 subunits assemble into a two-layered core harboring the HEPN catalytic centers, whereas the C-terminal domains form protrusions responsible for binding the ssDNA activator. We propose that DARNA remains inactive in uninfected cells due to the absence of accessible ssDNA (Figure 5C). The ssDNA generated during cellular processes such as DNA replication is bound by the bacterial SSB, which either sequesters the ssDNA or does not bind and present the ssDNA in a manner that can activate DARNA. Upon infection, phage SSB binds ssDNA fragments, likely including those produced during phage DNA replication, and this complex of phage SSB-ssDNA can potently activate DARNA.

Selective activation of DARNA by ssDNA bound to phage SSB, but not by ssDNA bound to bacterial SSB, implies that the accessibility of ssDNA to DARNA depends on whether it is associated with host or viral SSB proteins. Notably, T7, T5, and T4 phages belong to distinct viral clades and encode widely different SSBs, both in sequence and structure (Figure S1A). Moreover, while the phage SSBs T3 Gp2.5, T4 Gp32, T5 D11, and T7 Gp2.5 share a conserved oligonucleotide/oligosaccharide-binding (OB) fold with bacterial SSB ^38,43,44^, the overall similarity of the monomeric structures, including the position of ssDNA within the binding clefts, is greater between EcSSB and T3/T7 SSB than between T3/T7 SSB and DARNA-activating T4 and T5 phage SSBs (Figure S1B). This suggests that the key distinction between bacterial and phage SSBs lies not in monomer architecture, but in their modes of assembly on ssDNA. Whereas EcSSB forms homotetrameric complexes that wrap ssDNA tightly around the globular core ^45^, T4 and T7 phage SSBs assemble on ssDNA into filament-like structures ^46,47^. Likely due to the presence of linker ssDNA segments within these structures, together with significant SSB dissociation rates ^48^, such assemblies may expose ssDNA fragments in a manner that is sufficient for DARNA activation. Moreover, suppression of secondary structure formation in ssDNA and alterations in ssDNA conformation upon phage SSB binding ^46^ may account for the stronger activation of DARNA observed in the presence of both ssDNA and phage SSB compared with ssDNA alone (Figure 3A).

Structural analysis revealed that DARNA-ssDNA interactions are primarily mediated by protein contacts with DNA bases, including stacking interactions and base-specific hydrogen bonds with ‘CT’ dinucleotide fragments (Figure 4E). The ability of the DARNA dodecamer, but not the dimer, to bind ssDNA (Figure 3A) and the requirement for multiple ‘CT’ motifs within the activator oligonucleotide suggest that, for activation, DARNA relies on simultaneous binding to several ‘CT’ motifs in a single ssDNA fragment via multiple C-terminal domains. This model is consistent with the spatial arrangement of the ssDNA-binding domains on the exterior of the DARNA dodecameric assembly. However, the functional significance of the ‘CT’-motif specificity in the context of antiphage defense remains unclear.

The HEPN catalytic centers, located at the primary dimerization interfaces of DARNA subunits (three per homohexameric layer), are oriented toward the exterior of the dodecamer. Notably, in the apo structure, as well as in structures of DARNA bound to ssDNA alone or to both ssDNA and RNA, access to all catalytic centers is occluded by a protein loop (aa 81-85). We propose that activation of DARNA involves a rearrangement of this loop, allowing the tRNA substrate to access the catalytic center(s). This active, ssDNA-bound conformation may be transient or represent only a minor population under our experimental conditions, which could explain its absence in cryo-EM reconstructions, which likely capture the most abundant ground-state inactive complexes. The precise mechanism of tRNA recognition and cleavage by DARNA thus remains to be elucidated.

SSB proteins are among the most common phage determinants conferring phage sensitivity to diverse bacterial antiviral defense systems ^49^, including Retron-Eco8, DRT9, Hachiman, and others ^49–51^. Recent studies have shown that phage SSB binds directly to Eco8 and alters its msDNA conformation, thereby unleashing ATP-dependent DNA degradation ^10,52^, but the precise activation mechanism of other SSB-sensing systems remains to be established. In this study we demonstrated that an SSB protein, identified as a phage-encoded activator of the DARNA defense system through phage escaper mutant screening, does not interact with the defense protein directly. Instead, it increases the availability and likely the presentation of the direct activator, ssDNA. Such an indirect activation mechanism, in which viral proteins modify the accessibility or abundance of other, specific molecular cues rather than binding the defense effector itself, may represent a broader principle of immune system activation in bacteria. This mode of sensing expands the range of molecular patterns that can be recognized by the cellular immune systems, and enables them to achieve broad protection profiles by detecting common DNA species that arise during infection by highly diverse viruses.

### Methods Gene cloning

Arabinose-inducible plasmid constructs encoding phage SSB proteins were cloned by PCR (all oligonucleotide sequences are listed in Table S1) and Gibson assembly into pBAD30 and verified by commercial whole-plasmid sequencing. Site-directed mutagenesis of pBAD-Gp2.5 was conducted using 60 bp complementary primers containing the desired mutation and centered on the site to be mutated. Kapa polymerase was used to amplify from the WT plasmid template for 18 cycles. The reaction was treated with DpnI and directly transformed into competent cells. Mutagenized constructs were verified by whole-plasmid sequencing.

DARNA (PD-T7-3) gene was cloned via Gibson assembly into a pETDuet-1 vector with a C-terminal (His)_6_ affinity tag. Site-directed mutagenesis was performed to create the pETDuet-1 vector carrying the mutant variants of DARNA (H122A, F368A, W406A, R98E, R98E+K201E+K204E and F6A+N33A+E347A). Genes encoding phage SSB proteins (T4 gp32, T7 gp2.5, gp2.5-S132P, gp2.5-Y158C) were cloned via Gibson assembly into pACYCDuet-1 vectors. Sanger sequencing was performed to verify that the sequences were correct.

### Phage plaque assays

50 µL of overnight bacterial culture were mixed with 3.5 mL molten LB + 0.5% agar and overlaid onto a petri dish containing LB + 1.2% agar and 30 µg/mL chloramphenicol. A ten-fold dilution series of phage stocks were spotted onto the overlay and the plates were incubated at 37°C overnight.

### Bacterial growth assays

Bacterial strains containing pBAD constructs were grown overnight in LB + 0.2% glucose with the relevant antibiotics. The following day, cultures were diluted in fresh media and grown to OD600 = 0.4-0.5. These logarithmic phase cultures were washed of glucose and resuspended in LB with 0.2% arabinose. Cells were diluted to OD 0.05 in LB + 0.2% arabinose and 1.2 mL were distributed into the wells of a 24-well plate. Plates were shaken and incubated at 37°C in a BioTek Synergy H1 microplate reader and OD600 measurements were taken every 10 minutes. For agar plate assays, overnight cultures were washed of glucose, diluted in a 10-fold dilution series and spotted onto plates containing 0.2% arabinose. Plates were incubated at 30°C overnight before imaging.

### RNA-sequencing

Bacterial cultures were grown in liquid culture overnight. The following day, liquid cultures were diluted, grown to OD_600_ 0.4 and infected with T7 phage at MOI 5. Infected cultures were incubated at 37°C. At 12 minutes post-infection, 1 mL infected cultures were added to 111 µL of a stop solution (5% 0.1M citrate-buffered phenol, pH 4.3 in absolute ethanol). Cells were pelleted and resuspended in 65°C Trizol and RNA was isolated using the Zymogen Direct-zol RNA miniprep kit. Samples were then treated with DNase and RNA was precipitated and resuspended in nuclease free water. RNA samples were then treated with 20 units T4 Polynucleotide Kinase (PNK) to generate terminal 3′-phosphates from any 2′,3′-cyclic phosphate ends generated from HEPN RNase activity ^53^. Ribosomal RNAs were depleted using the method of Culviner et al. 2020 ^54^. RNA was then fragmented using the NEBNext Magnesium RNA Fragmentation Module. After precipitation, RNA sequencing libraries were generated using the Lexogen Small RNA-seq Library Prep Kit. Library quality and size distribution was confirmed using a 5300 Fragment Analyzer before 75 nt paired-end sequencing on an Illumina NextSeq500. Sequencing and demultiplexing was performed by the MIT BioMicro Center. Adaptor sequences were trimmed from reads using BBduk (ktrim = r k = 20 mink = 10 hdist = 1 tpe tbo, minlen=17). Reads were aligned to the MG1655 and T7 genomes using bowtie2 with default parameters. Read end and coverage analyses were performed using custom-made Python scripts employing Samtools as previously described ^23^.

### Gene expression and protein purification

All genes for protein purification were overexpressed in *E. coli* BL21(DE3) cells. Cultures carrying a gene of interest were grown at 37 °C in LB medium supplemented with the appropriate antibiotic (100 μg/ml of ampicillin or 30 μg/ml of chloramphenicol) until OD_600_ reached 0.5–0.7. Gene expression was then induced with 0.1 mM isopropyl β-D-1-thiogalactopyranoside. After induction, cells expressing DARNA gene (or its variants) were further grown for 4 hours at 37 °C, and cells expressing phage SSB genes were grown for 18 hours at 16 °C. Cells were collected by centrifugation and resuspended in lysis buffer (20 mM Tris-HCl (pH 8.0 at RT), 5 mM β-mercaptoethanol (2-ME), 2 mM phenylmethylsulfonyl fluoride), which also contained 500 mM NaCl for DARNA and 50 mM NaCl for phage SSB proteins. Cells were lysed by sonication, cell debris was removed by centrifugation and the supernatant was collected for protein purification. For DARNA (and its variants) purification, the samples were loaded on a Ni^2+^-charged HiTrap chelating HP column (Cytiva) and the protein of interest was eluted using a linear gradient (5%-100%) of an imidazole containing buffer (20 mM Tris-HCl (pH 8.0 at RT), 500 mM NaCl, 500 mM imidazole, 5 mM 2-ME and 5 % (v/v) glycerol). Fractions containing DARNA protein were pooled and diluted 2-fold with 20 mM Tris-HCl (pH 8.0 at RT) buffer, then loaded on a HiTrap heparin HP column (Cytiva). Protein was eluted using a linear gradient (25%-100%) of a high-salt buffer (20 mM Tris-HCl (pH 8.0 at RT), 1M NaCl, 5 mM 2-ME and 5 % (v/v) glycerol). Phage SSB proteins containing samples were initially loaded on a HiTrap heparin HP column and the proteins were eluted using a linear gradient (10%-100%) of a high-salt buffer. Fractions containing an SSB protein were pooled and diluted to 50 mM NaCl concentrations with 20 mM Tris-HCl (pH 8.0 at RT) buffer, then loaded on a HiTrap Q HP column (Cytiva). Protein was eluted using a linear gradient (10%-100%) of a high-salt buffer. All purified proteins were dialyzed against a storage buffer (20 mM Tris-HCl (pH 8.0 at RT), 500 mM NaCl, 2 mM dithiothreitol (DTT) and 50% (v/v) glycerol) and checked on a 12 % SDS-PAGE.

### tRNA cleavage reactions in vitro

For tRNA cleavage in vitro, 3 μM of *E. coli* tRNAs was mixed with 0.5 μM (monomer concentration) of DARNA, 0.1 μM of ssDNA and 5 μM of phage SSB protein in the reaction buffer (60 mM Tris-acetate (pH 8.0 at RT), 132 mM potassium acetate). Samples were incubated at 37 °C for 5 min, (reaction time was extended to 20 min when *E. coli* SSB was also added), reactions were stopped by incubating samples at 80 °C for 10 min. Afterwards, the samples were diluted 3-fold, treated sequentially with DNase I and proteinase K, and then mixed with 2× RNA loading dye. The samples were analyzed by 10% urea (8 M) polyacrylamide gel electrophoresis (PAGE) in 1× TBE buffer (Thermo Scientific), stained with SYBR Gold, and visualized using an Amersham Typhoon biomolecular imager.

### Mass Photometry

Mass photometry measurements were performed using a Refeyn TwoMP instrument. Prior to measurements, the instrument was calibrated using a protein standard mix (MassFerence P1). Sample components were diluted in buffer (60 mM Tris-acetate (pH 8.0 at RT), 132 mM potassium acetate) to the following concentrations: 100 nM (monomer) DARNA, 50 nM ssDNA, 150 nM phage SSB. 10 µL of buffer was added for focusing. Subsequently, 10 µL of the sample was introduced, and movies were recorded for 60 s at room temperature. Data were analyzed using Refeyn Discover software. Mass histograms were fitted with Gaussian functions to estimate the distribution and relative abundance of different molecular species.

### Cryo-EM sample preparation

For cryo-EM, DARNA was dialyzed against a buffer containing 50 mM Tris-acetate (pH 7.5), 150 mM potassium acetate, and 1 mM dithiothreitol (DTT). 3 µl of the sample (12 µM DARNA monomer) was applied to a glow-discharged (20 mA for 45 s) Quantifoil Cu grids (300 mesh, R 1.2/1.3) and vitrified in liquefied ethane using Vitrobot Mark IV (FEI) at 4 °C and 100% humidity with 0 s waiting and 6 s blotting times. For samples containing ssDNA, components were mixed directly on the grid, final concentrations were 9 µM DARNA-H122A and 9 µM ssDNA (34 nt). For samples containing ssDNA and *E. coli* tRNA mix, solution of ssDNA and tRNA was prepared beforehand and mixed with DARNA-H122A directly on the grid, final concentrations were 15 µM DARNA-H122A 15 µM ssDNA (34 nt) and 50 µM tRNA mix.

### Cryo-EM data collection and image processing

Cryo-EM data collection is summarized in Table S2. Data collection was performed using a Glacios microscope (Thermo Fisher Scientific), operating at 200 kV and equipped with a Falcon 3EC Direct Electron Detector in the electron counting mode (Vilnius University, Vilnius, Lithuania). Data was collected with EPU (v.3.2 - v.3.13) at a nominal magnification of 92,000×, corresponding to a calibrated pixel size of 1.10 Å, using an exposure of 0.80 e^-^/Å^2^ s^−1^, in 30 frames and a final dose of 30 e^-^/Å^2^, over a defocus range of −1.0 to −2.0 µm. Patch motion correction, CTF estimation, micrograph curation and, in some cases, blob picking with particle extraction were performed in real-time in CryoSPARC Live (v.4.1.0 - v.4.7.1) ^55^. Further data processing and final refinement were performed using standard CryoSPARC (v.4.1.0 - v.4.7.1) ^55,56^. Image processing for each sample is summarized in Figures S7-S9. In all cases, 3DFSC web server ^57^ was used to estimate the global resolution at 0.143 FSC cutoff and sphericity values of the final electron density maps, while the local resolution was calculated using CryoSPARC (v.4.7.1).

### Cryo-EM model building, refinement and analysis

The initial model of the DARNA monomer was generated using AlphaFold 3 ^58^ and twelve models were fitted into the cryo-EM map of the dodecameric complex using ChimeraX (v.1.9)^59^. Protein rebuilding was performed using Coot (v.0.9.8.1) ^60^. For building structures with ssDNA and tRNA, the initial model of the dodecamer in the apo state was used. Models were adjusted and fragments of ssDNA and RNA were manually built using Coot (v.0.9.8.1) ^60^. Models were refined using phenix.real_space_refine (v. 1.21.2-5419) ^61^, refinement statistics are summarized in Table S2. Structure overlays and generation of structural images were performed using ChimeraX (v.1.9) ^59^.

### Electrophoretic mobility shift assay (EMSA)

A ssDNA oligonucleotide (34 nt) was radiolabeled at 5′ end with [γ-³²P]-ATP using T4 polynucleotide kinase. The oligonucleotide (98 nM unlabeled + 2 nM radiolabeled) was diluted in the binding buffer (40 mM Tris-acetate pH 8.3 (RT), 100 mM potassium acetate, 10 (v/v) % glycerol, and 0.1 mg/ml BSA) and mixed with increasing concentrations of SSB protein (0-5 μM). Samples were incubated at RT for 10 minutes and analyzed by running a 8% PAGE in a 40 mM Tris-acetate (pH 8.3 at RT) buffer. Results were visualized by scanning an autoradiography phosphor screen using an Amersham Typhoon biomolecular imager.

## DATA AVAILABILITY

The EM densities of the DARNA structures (composite map, consensus map, focused map 1 obtained with a mask covering the C-terminal parts of A-B subunits, and focused map 2 obtained with a mask covering the C-terminal part of subunit C) have been deposited in the Electron Microscopy Data Bank under the accession codes EMD-56164/EMD-56161/EMD-56162/EMD-56163 (DARNA in apo-form), EMD-56168/EMD-56165/EMD-56166/EMD-56167 (DARNA in complex with ssDNA), EMD-56172/EMD-56169/EMD-56170/EMD-56171 (DARNA in complex with ssDNA and tRNA), and EMD-5676/EMD-56173/EMD-56174/EMD-56175 (DARNA in complex with tRNA). The DARNA model coordinates have been deposited in the Protein Data Bank under the accession codes 9tra (apo-form), 9trc (complex with ssDNA), 9trd (complex with ssDNA and tRNA), and 9tre (complex with tRNA).

## ACKNOWLEDGEMENTS

We thank Darius Kazlauskas for helpful discussions on SSB proteins.

## Supplementary Material

**Figure S1.**
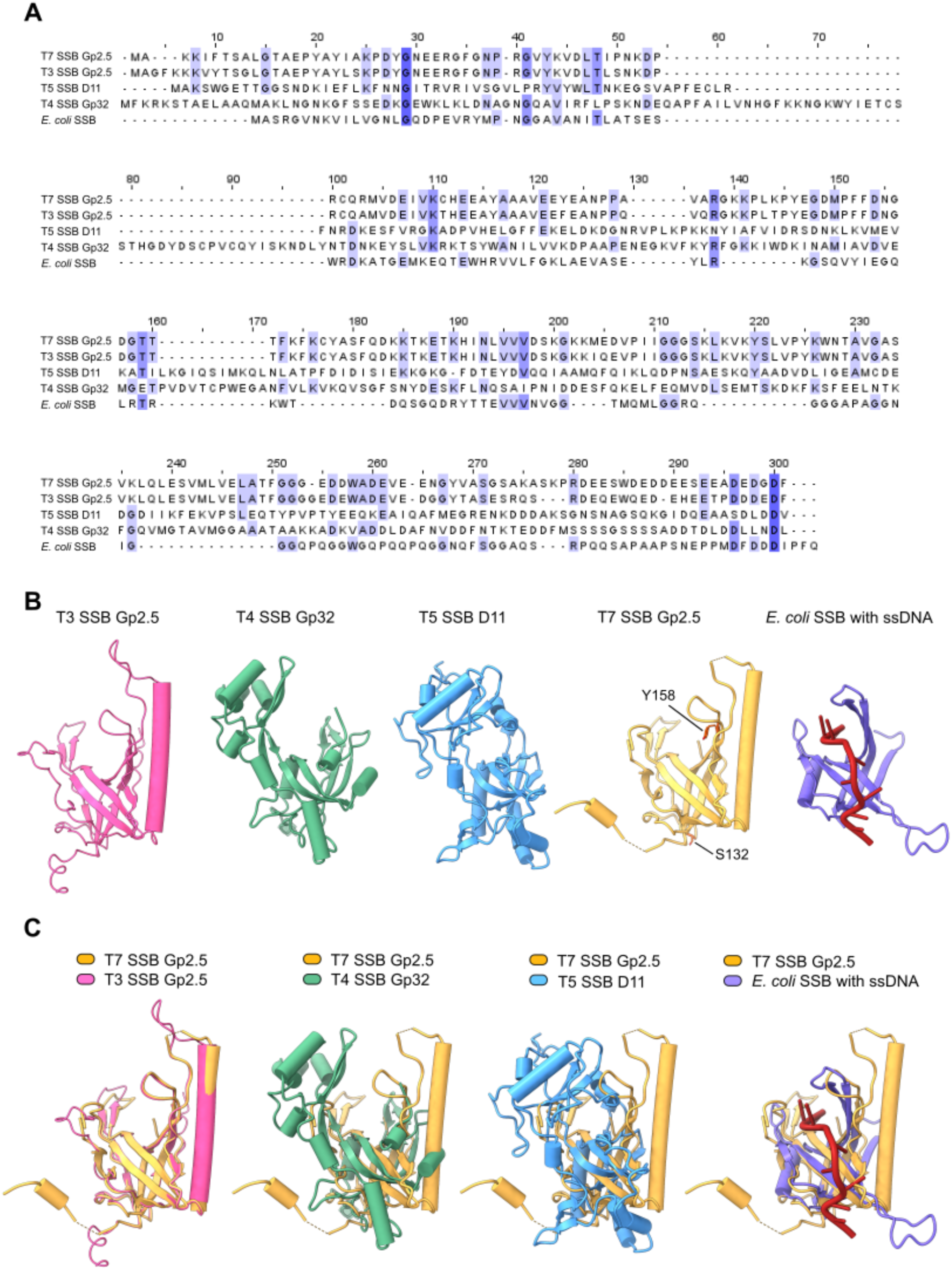
Comparison of single-stranded DNA-binding proteins. (A) Multiple sequence alignment of different phage and *E. coli* SSB proteins. Residues are colored according to percentage identity. (B) Structures of different SSB proteins. Left to right: phage T3 SSB Gp2.5 (AlphaFold 3 ^58^ model), phage T4 SSB Gp32 (PDB ID 1gpc), phage T5 SSB D11 (AlphaFold 3 ^58^ model), phage T7 SSB Gp2.5 (PDB ID 1je5) and residues that are critical for activation of DARNA, *E. coli* SSB with bound ssDNA (PDB ID: 1eyg, only one protein subunit shown). (C) Left to right: overlay of T7 SSB Gp2.5 (PDB ID 1je5) and T3 SSB Gp2.5 (AlphaFold 3 ^58^ model), overlay of T7 SSB Gp2.5 (PDB ID 1je5) and T4 SSB Gp32 (PDB ID 1gpc), overlay of T7 SSB Gp2.5 (PDB ID 1je5) and T5 SSB D11 (AlphaFold 3 ^58^ model), overlay of T7 SSB Gp2.5 (PDB ID 1je5) and *E. coli* SSB monomer with bound ssDNA (PDB ID 1eyg).

**Figure S2.**
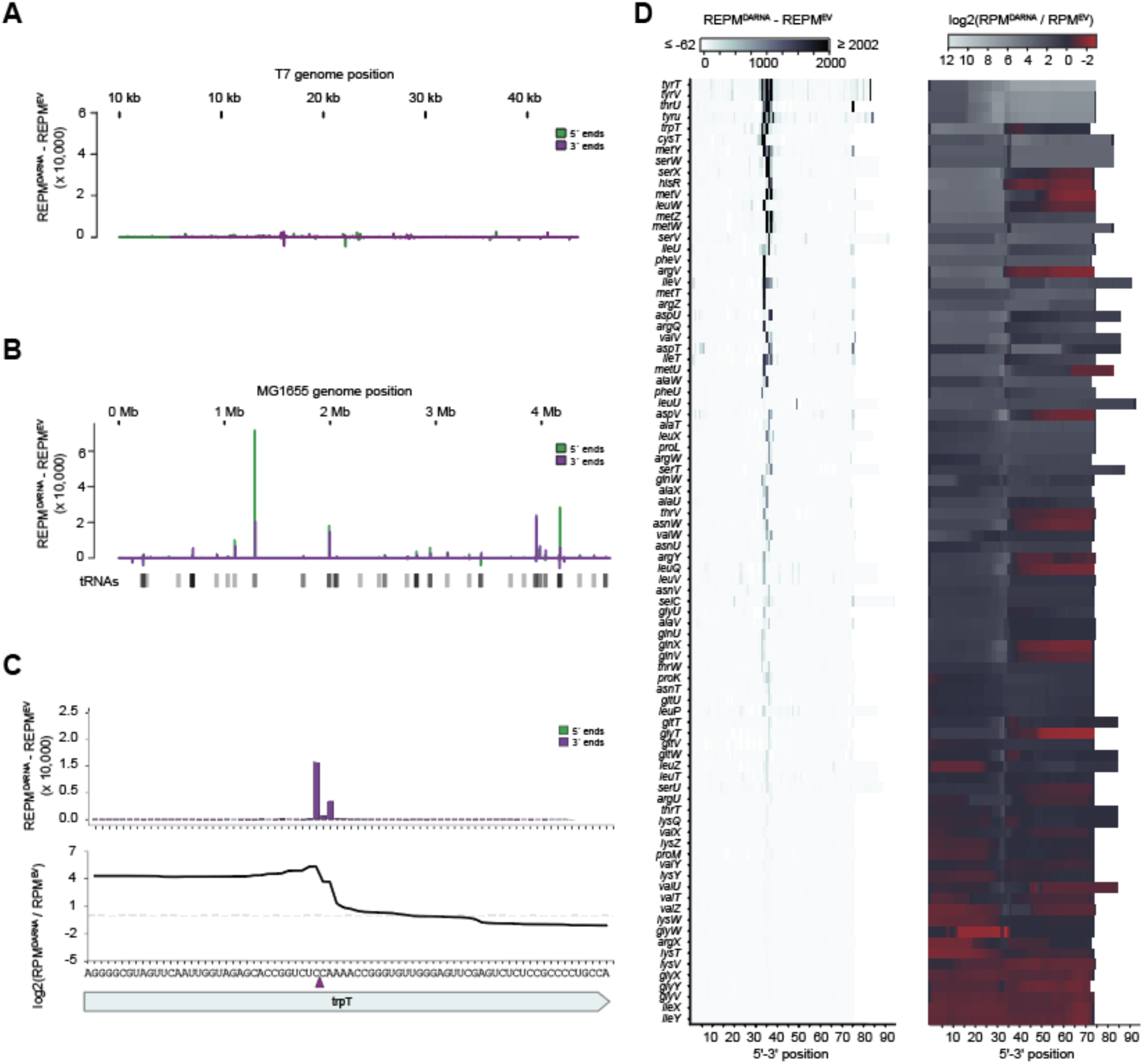
Identification of target RNAs cleaved by DARNA. (A) RNA-sequencing data of T7 infected empty vector cells or DARNA-encoding cells expressed as normalized read ends of the DARNA strain subtracted from the normalized read ends of the EV strain at each position along the T7 chromosome. (B) RNA-sequencing data measuring read ends from uninfected DARNA cells expressing Gp2.5 S132P relative to those expressing Gp2.5. (C) Same data as (B), but enlarged to show the *trpT* tryptophan tRNA gene (top). Bottom: coverage ratio at positions along the gene. Red arrow indicates presumed cleavage site. (D) Left, heatmap showing the same metric as (B) but within all canonical tRNAs. To better show detail, the colorbar maximum and minimum are at the 99.5 and 0.05 percentile of the total data, respectively. Right, coverage ratio at positions along tRNAs.

**Figure S3.**
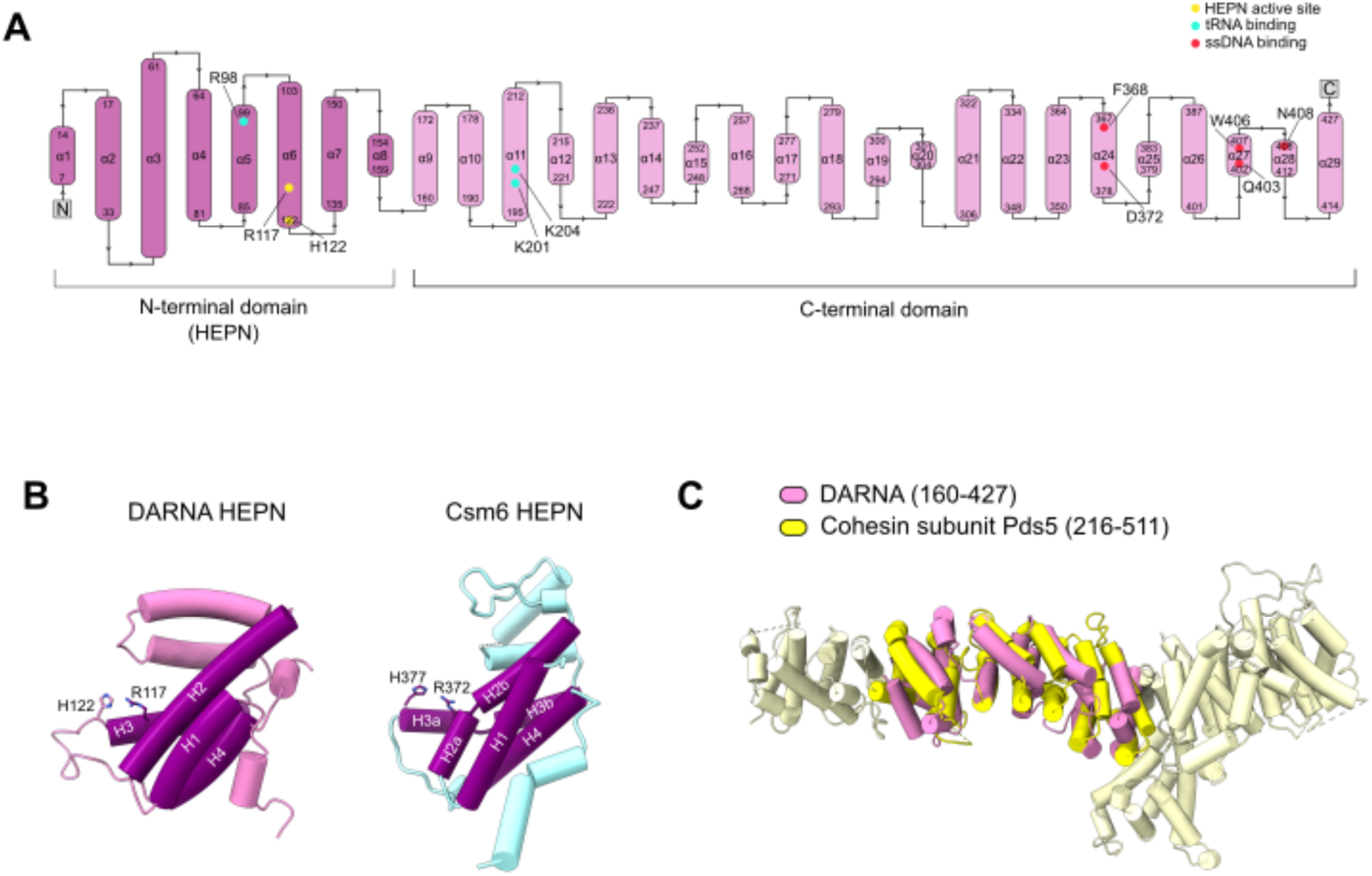
Structural features of DARNA. (A) Topology diagram of DARNA. The HEPN catalytic center, tRNA binding residues, and ssDNA binding residues are highlighted. (B) Comparison of the HEPN domains of DARNA and Csm6 (PDB ID: 6tug). The DALI ^41^ Z score for the overlay is 8.3 (RMSD 2.9 over 116 Cα atoms). Canonical HEPN α-helices are highlighted in purple and marked H1-H4 as described in ^62^. HEPN active site residues (H122/R117 and H377/R372 for DARNA/Csm6 respectively) are shown in stick representation. (C) Comparison of DARNA C-terminal domain with the cohesin subunit Pds5 (PDB ID: 5f0o). The overlay was obtained using GTalign ^42^, yielding TM score of 0.6387 and an RMSD of 5.17, with 272 residues aligned.

**Figure S4.**
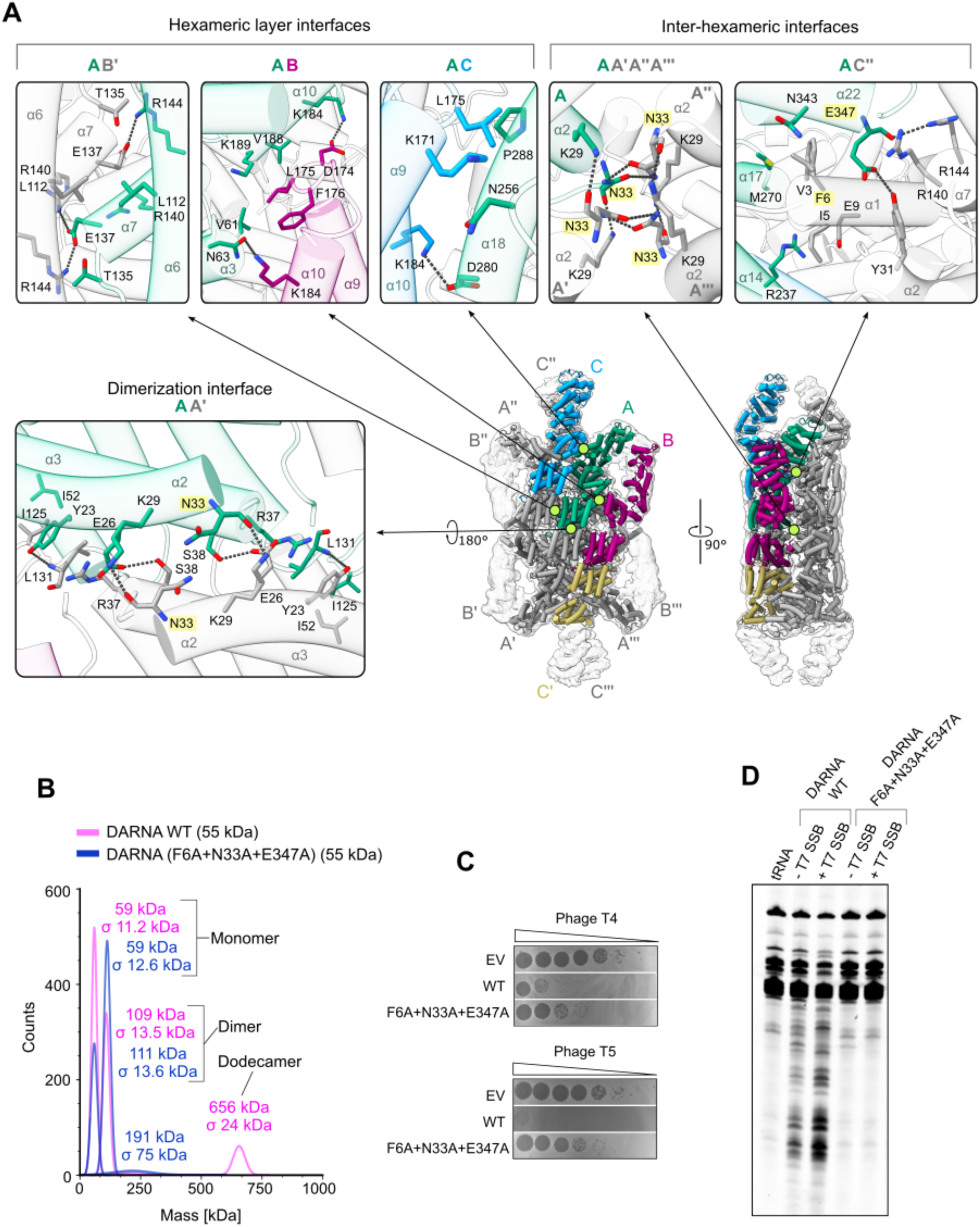
Oligomerization interfaces of DARNA. (A) Light green circles indicate the subunit interfaces within the dodecameric assembly that are shown in the corresponding inset panels. Residues in the zoom-in views are colored according to the subunit coloring used in the overall DARNA structure. Substituted residues are highlighted in yellow. (B) Mass photometry measurements showing that F6A+N33A+E347A DARNA variant (blue) predominantly exists as a monomer or a dimer, while WT (magenta) can also form a dodecamer. Mass photometry data for the WT protein is also shown in Fig. 3B. (C) Phage plaque assays with phages T4 and T5 comparing DARNA WT and F6A+N33A+E347A variants. (D) tRNA cleavage assays comparing DARNA WT and F6A+N33A+E347A variants. tRNA cleavage reactions contained DARNA and T7 SSB at concentrations of 0.5 μM and 5 μM, respectively (in terms of monomer), and ssDNA at 0.1 μM.

**Figure S5.**
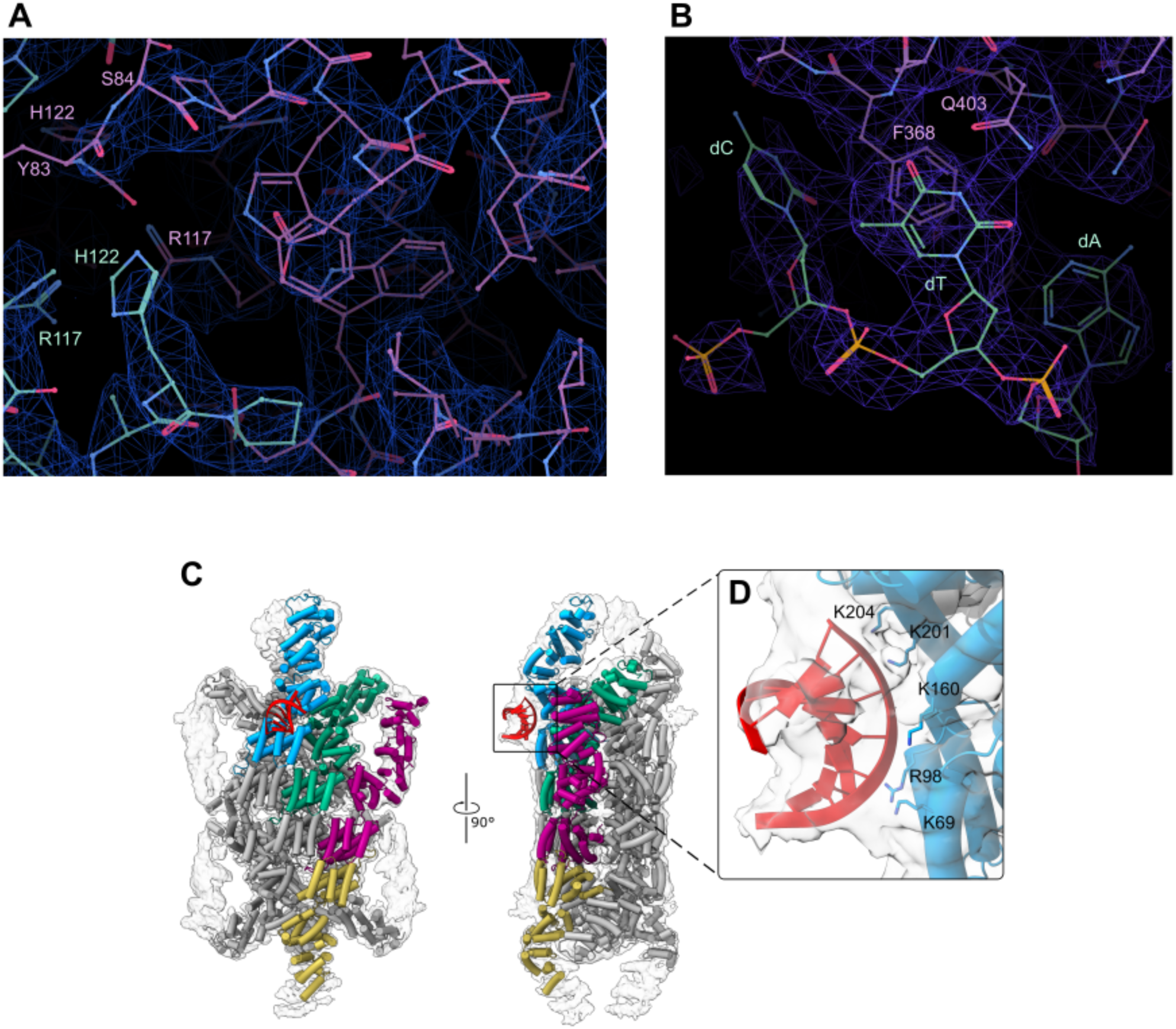
Cryo-EM studies of DARNA in apo-and nucleic acid-bound forms. (A) Sharpened cryo-EM volume and atomic model covering the HEPN catalytic center region in the apo-DARNA structure. (B) Sharpened cryo-EM volume and atomic model of ssDNA fragment ‘CTA’ bound to DARNA. (C) Overall structure of the DARNA homododecamer bound to an RNA fragment in the absence of ssDNA. The unsharpened cryo-EM map (transparent) and the atomic model are shown (D) DARNA residues contacting the bound RNA in the DARNA-RNA complex.

**Figure S6.**
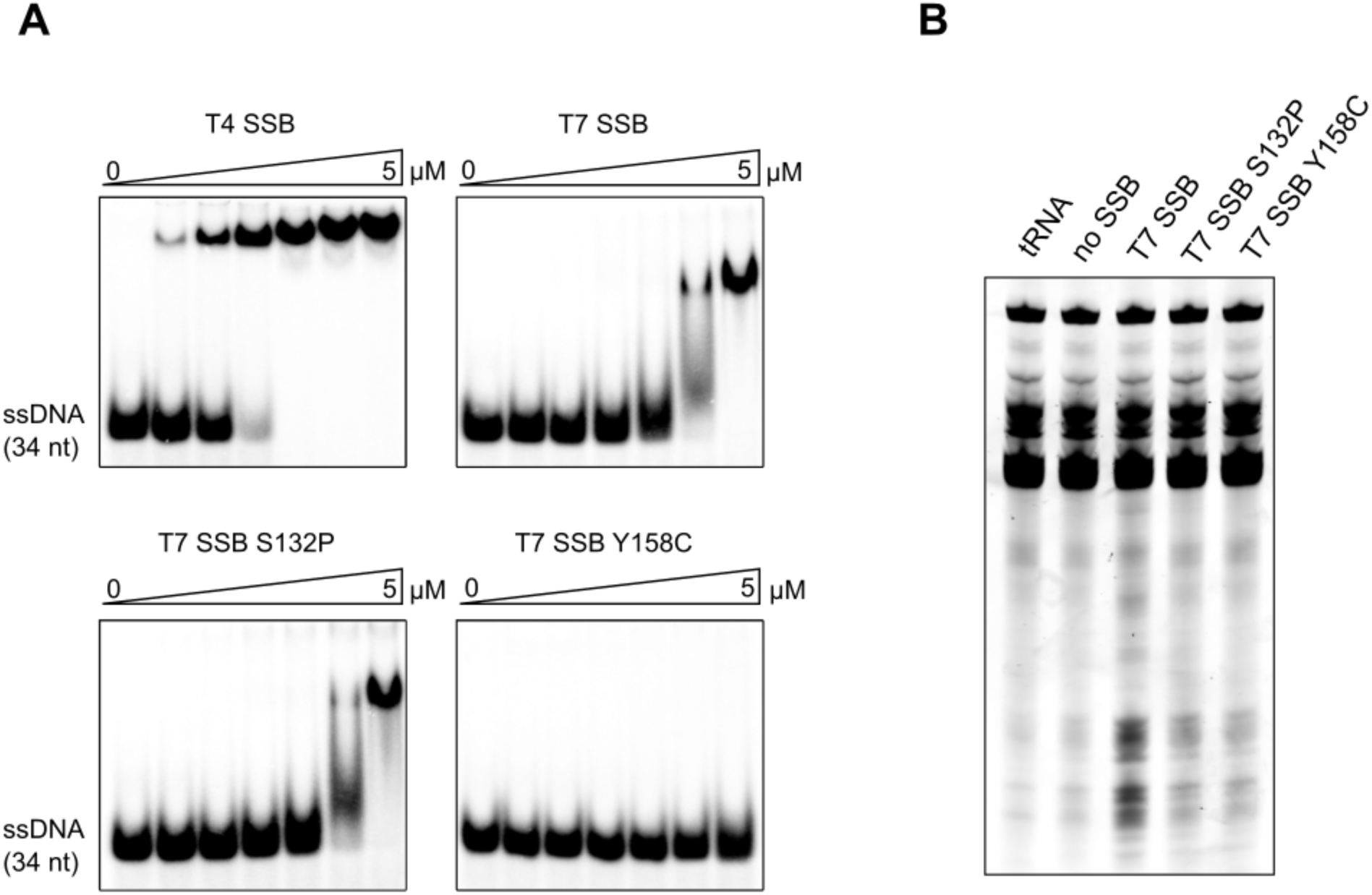
SSB proteins in DARNA reactions. (B) EMSA using different phage SSB variants (T4 SSB, T7 SSB, T7 SSB S132P, T7 SSB Y158C) and ‘34nt’ ssDNA. ssDNA concentration is 100 nM, protein concentrations are 0, 100, 200, 500, 1000, 2500, 5000 nM. (C) tRNA cleavage assays testing the ability of T7 SSB variants (5 µM) to outcompete *E. coli* SSB protein (0.8 µM) and activate DARNA.

**Figure S7.**
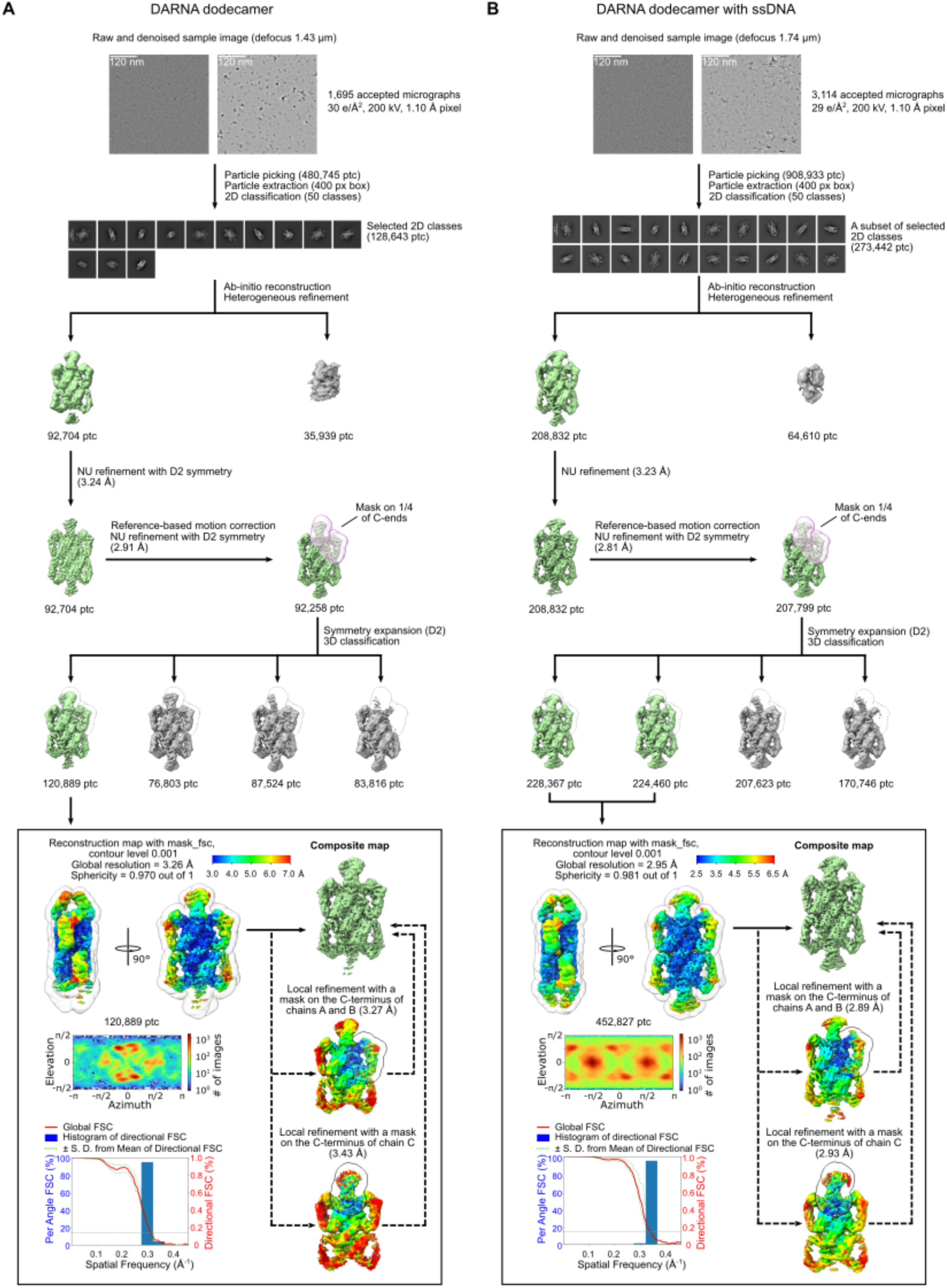
Cryo-EM data processing. (A) apo-DARNA. (B) DARNA with bound ssDNA.

**Figure S8.**
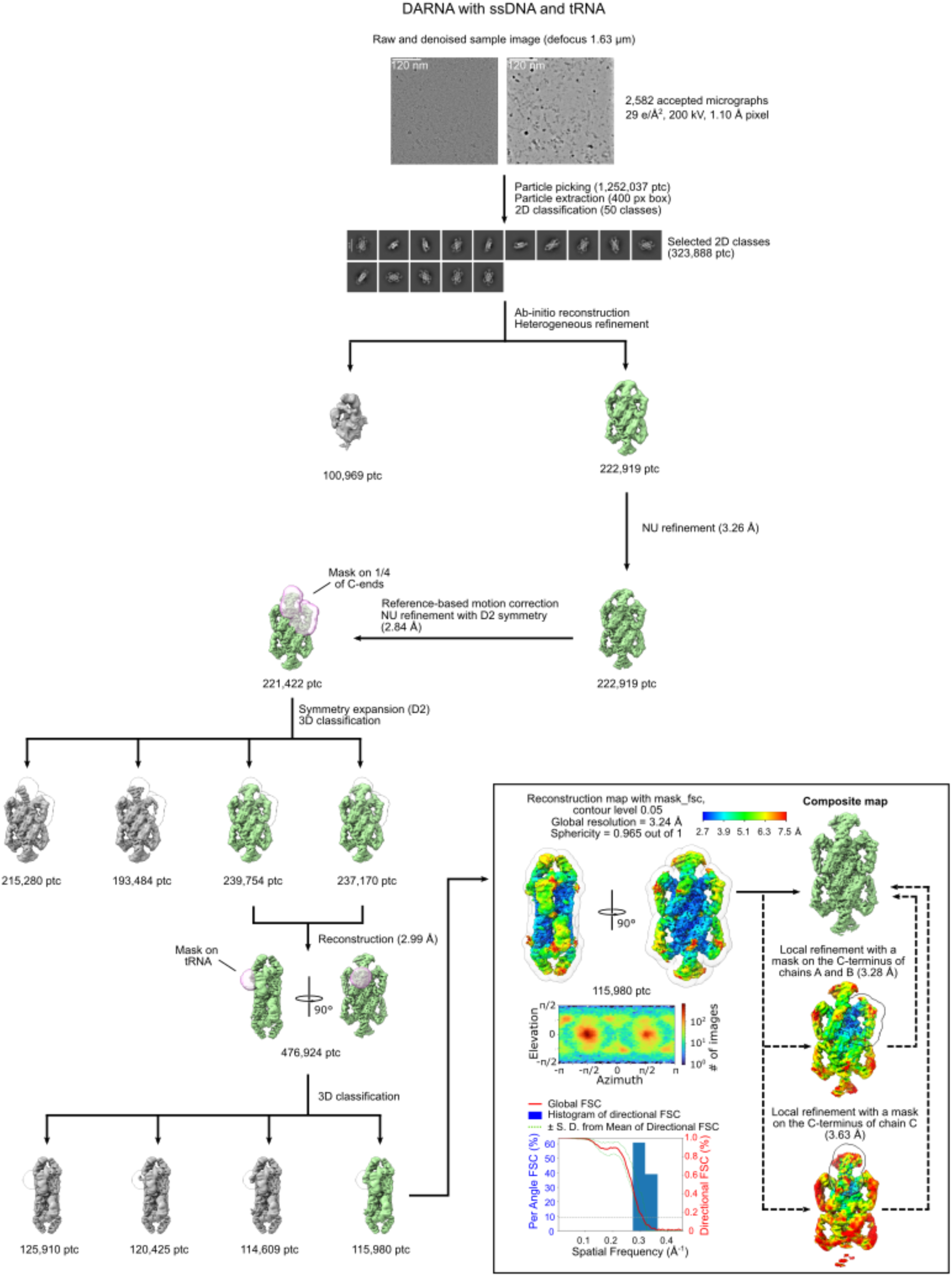
Cryo-EM data processing for DARNA with bound ssDNA and tRNA.

**Figure S9.**
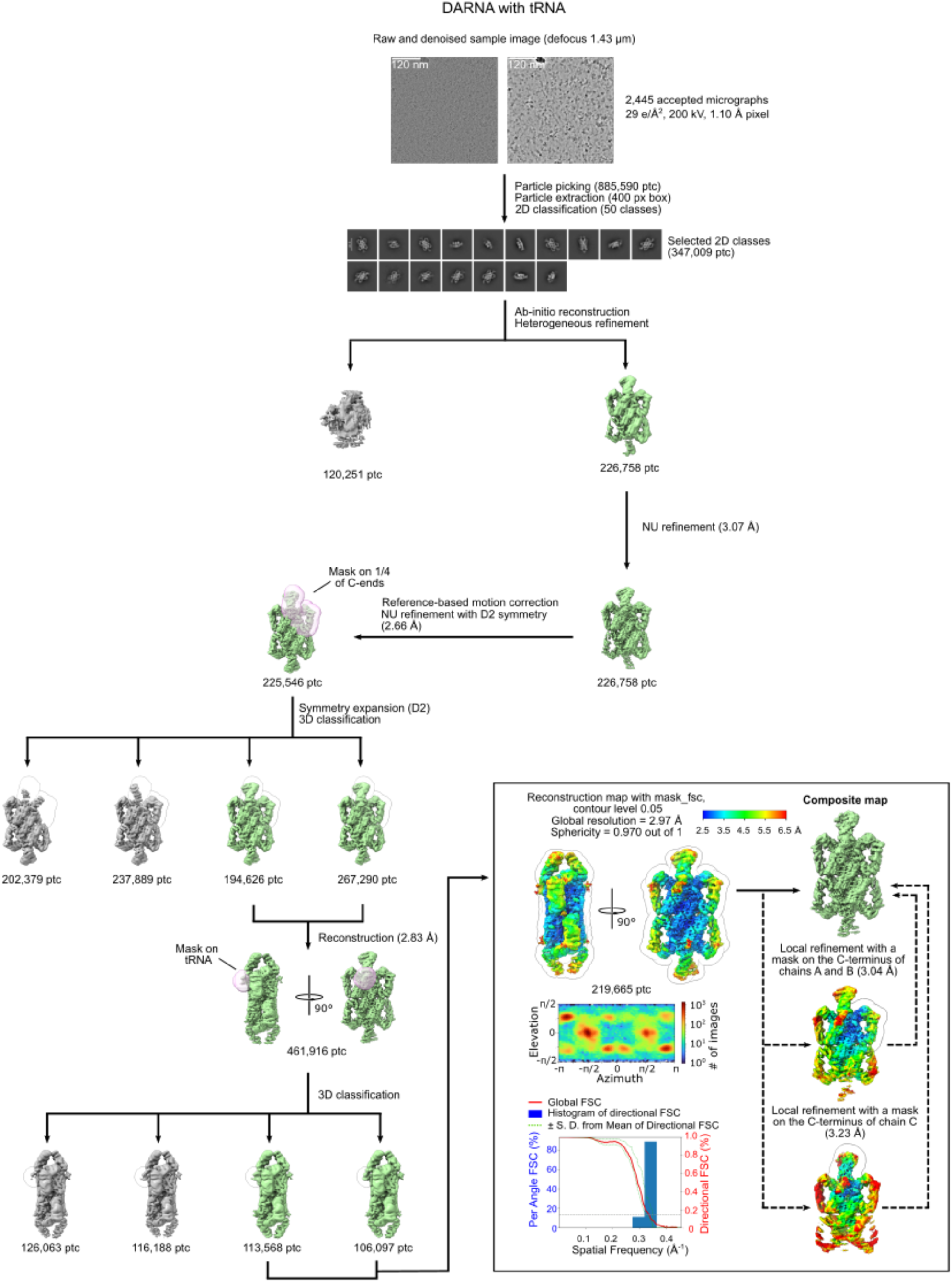
**Cryo-EM data processing for DARNA with bound tRNA.**

---

**Table S1.**
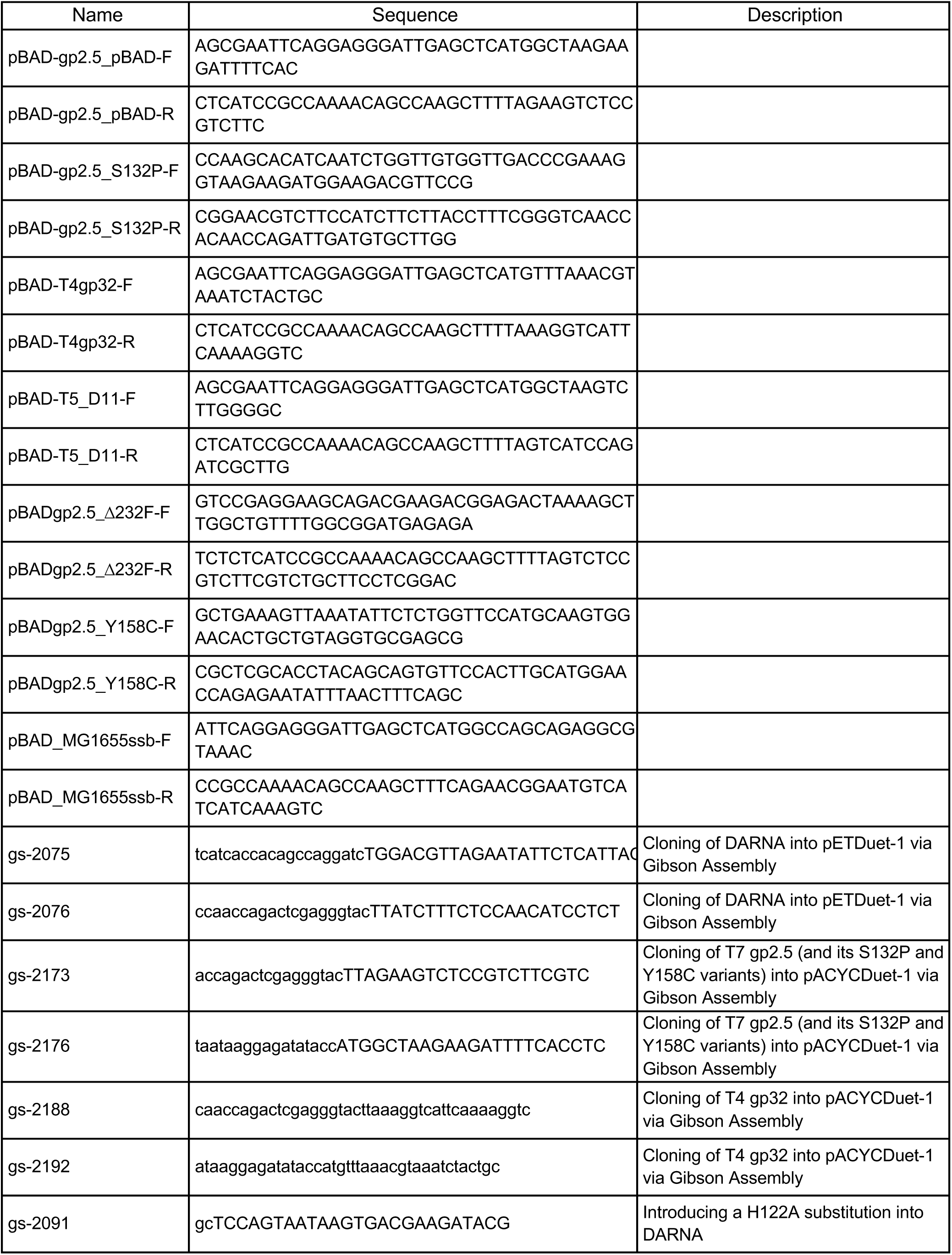
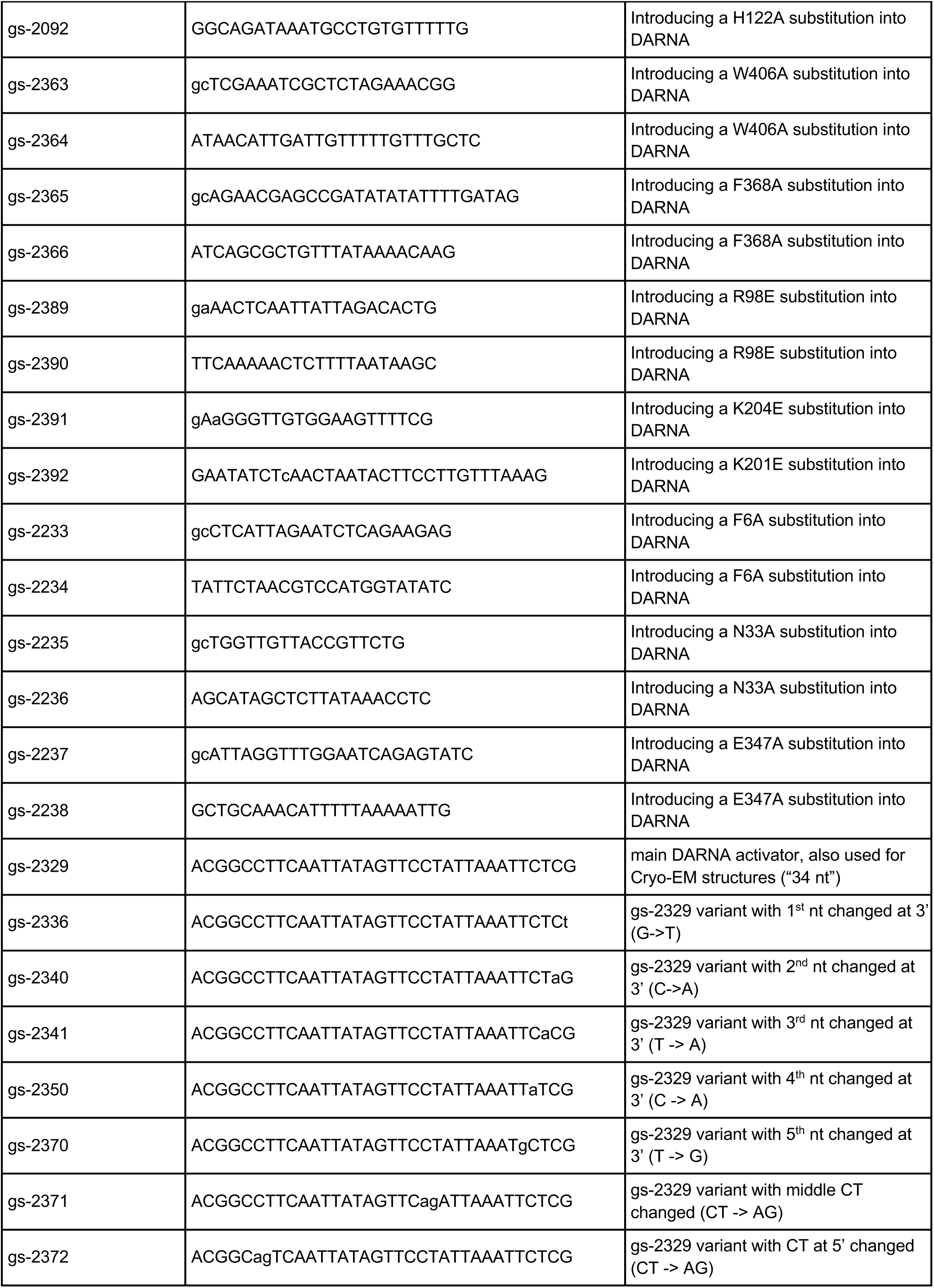
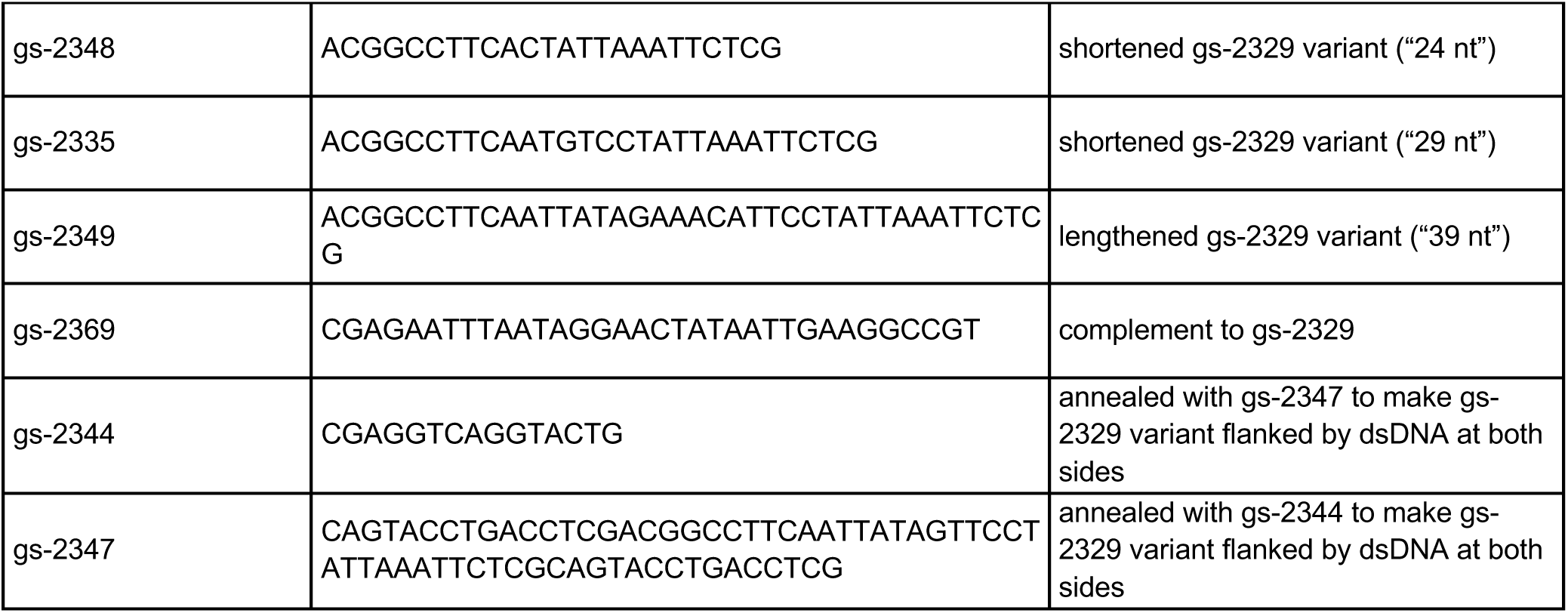
DNA oligonucleotides.

**Table S2.** Cryo-EM data collection, refinement and validation statistics.

|  | DARNA | DARNA + ssDNA | DARNA + ssDNA + tRNA | DARNA + tRNA |
| --- | --- | --- | --- | --- |
| PDB | 9TRA | 9TRC | 9TRD | 9TRE |
| EMDB* | EMD-56164<br>EMD-56161<br>EMD-56162<br>EMD-56163 | EMD-56168<br>EMD-56165<br>EMD-56166<br>EMD-56167 | EMD-56172<br>EMD-56169<br>EMD-56170<br>EMD-56171 | EMD-56176<br>EMD-56173<br>EMD-56174<br>EMD-56175 |
| Data collection and processing |  |  |  |  |
| Microscope | Glacios (TFS) | Glacios (TFS) | Glacios (TFS) | Glacios (TFS) |
| Detector | Falcon III camera (TFS) | Falcon III camera (TFS) | Falcon III camera (TFS) | Falcon III camera (TFS) |
| Magnification | 92,000 | 92,000 | 92,000 | 92,000 |
| Automation software | EPU | EPU | EPU | EPU |
| Voltage (kV) | 200 | 200 | 200 | 200 |
| Electron exposure (e<sup>-</sup>/Å<sup>2</sup>) | 30.0 | 29.0 | 29.0 | 29.0 |
| Defocus range (μm) | 0.5–2.5 | 0.5–2.5 | 0.5–2.5 | 0.5–2.5 |
| Pixel size (Å) | 1.1 | 1.1 | 1.1 | 1.1 |
| Micrographs collected/used | 1,822/1,695 | 3,727/3,114 | 2,810/2,582 | 2,804/2,445 |
| Initial/final particle images (no.) | 480,745/120,889 (after D2 symmetry expansion) | 908,933/452,827 (after D2 symmetry expansion) | 1,252,037/115,980 (after D2 symmetry expansion) | 885,590/219,665 (after D2 symmetry expansion) |
| Symmetry imposed | C1 | C1 | C1 | C1 |
| Map resolution (Å) (masked/unmasked); FSC threshold 0.143 | 3.26/4.36* | 2.95/3.73* | 3.24/4.35* | 2.97/3.97* |
| Map sharpening B factor ($\text{\AA}^2$) | 107.88* | 120.15* | 86.47* | 70.28* |
| 3D FSC sphericity score | 0.970* | 0.981* | 0.965* | 0.970* |
| Refinement (Phenix) |  |  |  |  |
| Initial model used | AlphaFold 3 | PDB ID: 9TRA | PDB ID: 9TRC | PDB ID: 9TRD |
| Model resolution ($\text{\AA}$)<br>FSC threshold 0.5 | 3.3 | 3.1 | 3.5 | 3.4 |
| Model composition |  |  |  |  |
| Non-hydrogen atoms | 33,892 | 35,659 | 35,742 | 34,430 |
| Protein residues | 4,277 | 4,484 | 4,477 | 4,353 |
| Nucleotides | 0 | 21 | 37 | 14 |
| B factors ($\text{\AA}^2$) min/max/mean | | | | |
| Protein | 25.25/218.88/105.19 | 91.05/382.27/180.36 | 47.56/381.16/159.01 | 8.25/356.30/141.03 |
| Nucleotide |  | 176.88/328.64/272.36 | 132.30/503.15/241.56 | 212.28/331.35/279.08 |
| RMSDs |  |  |  |  |
| Bond lengths ($\text{\AA}$) | 0.002 | 0.003 | 0.002 | 0.002 |
| Bond angles ($^\circ$) | 0.417 | 0.462 | 0.490 | 0.451 |
| Validation |  |  |  |  |
| MolProbity score | 1.30 | 1.34 | 1.39 | 1.52 |
| Clashscore | 5.60 | 5.66 | 7.17 | 8.04 |
| Poor rotamers (%) | 0.54 | 1.10 | 0.78 | 0.20 |
| Ramachandran plot |  |  |  |  |
| Favored (%) | 98.47 | 98.14 | 98.27 | 97.60 |
| Allowed (%) | 1.53 | 1.86 | 1.73 | 2.40 |
| Disallowed (%) | 0 | 0 | 0 | 0 |
| CCvolume/CCmask | 0.79/0.79 | 0.83/0.83 | 0.85/0.85 | 0.84/0.85 |
| Cβ outliers (%) | 0 | 0 | 0 | 0 |
| CaBLAM outliers (%) | 0.47 | 0.52 | 0.54 | 0.65 |

\*Four EMDB codes are shown: the composite map, the consensus map, focused map 1 (mask covering C-terminal parts of subunits A and B), and focused map 2 (mask covering the C-terminal part of subunit C); map resolution and sphericity scores are reported for the consensus map; map sharpening B factor is reported for the composite map.

